# Higher-order assembly of a type IX retron enables exploitation for designer antimicrobials

**DOI:** 10.64898/2026.07.11.737809

**Authors:** Grace N. Hibshman, Linhan Wang, Nicole MacRae, Karen Zhang, Alfredo Florez, Seth L. Shipman, Eva Nogales

**Affiliations:** California Institute for Quantitative Biosciences and Department of Molecular and Cell Biology, University of California, Berkeley, CA, USA; Howard Hughes Medical Institute, University of California, Berkeley, Berkeley, CA, USA; Gladstone Institute of Data Science and Biology, San Francisco, CA, USA; Biophysics Graduate Group, University of California, Berkeley, Berkeley, CA, USA; Department of Bioengineering and Therapeutic Sciences, University of California, San Francisco, CA, USA; Chan Zuckerberg Biohub, San Francisco, CA, USA; Department of Molecular and Cell Biology, University of California, Berkeley, Berkeley, CA, USA; Molecular Biophysics and Integrated Bioimaging Division, Lawrence Berkeley National Laboratory, Berkeley, CA, USA

## Abstract

Bacterial defense systems provide a rich reservoir for biotechnological innovation. Retrons are tripartite abortive infection systems that detect phage invasion using reverse-transcribed DNA (msDNA), but how they structurally couple threat detection to effector activation remains poorly understood. Here, we determine the cryo-EM structure and activation mechanism of retron-Kva2, a type IX retron from the human pathogen *Klebsiella variicola*. We reveal that retron-Kva2 assembles into an asymmetric, higher-order ribonucleoprotein complex that sequesters a toxic dimeric HEPN RNase at its core. We identify a natural phage trigger as the phage T5 protein D5, which activates the retron through structural mimicry. Mirroring the retron-Kva2 winged-helix protein, the helix-turn-helix fold of D5 binds the msDNA sensor, driving conformational remodeling that unleashes HEPN-mediated tRNA cleavage and growth arrest. Because retron-Kva2 surveils a structural fold via msDNA binding, rather than a primary sequence, this recognition mechanism provides a broadly exploitable pathway for programmable activation. Harnessing this structure-based logic, we computationally designed *de novo* synthetic triggers that activate retron-Kva2-mediated bacterial growth arrest *in vivo*. Our findings reveal the architectural basis of type IX retron immunity and establish a structure-guided paradigm for repurposing bacterial defense systems into precision-honed antimicrobial therapeutics.

## Introduction

Bacteria have been under threat from bacteriophage and other selfish genetic elements for billions of years. In response, bacteria have evolved a remarkable arsenal of immune systems to defend against these invaders. Bacterial immune systems have also been co-opted by researchers to create molecular tools. Restriction modification and CRISPR-Cas systems are two examples of bacterial immune systems that have been transformed into revolutionary technologies with wide-ranging implications, from research to therapeutics^1,2^. Recent systematic analyses of microbial genomes have uncovered thousands of new defense systems with entirely uncharacterized mechanisms that hold potential for development into novel biotechnology^3–7^. One category of defense system with outsized biotechnological potential are retrons, which consists of three main components: a non-coding RNA (ncRNA) with two parts, the multicopy single-stranded RNA (msrRNA) and the multicopy single-stranded DNA-coding RNA (msdRNA) which serves as a template for reverse transcription, a reverse transcriptase (RT) that copies the msdRNA into msdDNA, generating a unique RNA-DNA chimera called msDNA, and an effector protein, which varies widely in predicted functions^8^.

Retrons were first discovered in the 1980s, yet their biological role remained unclear until 2020, when they were classified as abortive infection (Abi) defense systems in which the RT-msrRNA-msdDNA module functions as the sensor for phage infection, and an associated effector protein halts the propagation of the phage by targeting bacterial host processes^4,8–11^. Retrons have since been grouped into sixteen subtypes distinguished by the effector identity, ranging from ATPases, cold-shock proteins, HNH endonucleases, HEPN ribonucleases (RNase), to transmembrane proteins, with more still being classified^12^. To date, only a few retrons have been thoroughly characterized, and we lack a mechanistic understanding of how most retron types sense and respond to infection. Although the retron RT and ncRNA have already been engineered into tools, including genome editing and molecular recording, the effector component that executes defense remains unexplored for biotechnology^13–16^.

Type IX retrons are particularly intriguing, as they encode two distinct effector proteins, a higher eukaryotes and prokaryotes nucleotide-binding (HEPN) protein and a winged-helix (WH) DNA-binding protein within the same operon as the RT and ncRNA^12^. HEPN domains are minimal RNases that act as homodimers^17^. They are deployed across kingdoms in immune and stress-response pathways, including CRISPR–Cas13, the type III CRISPR effector Csm6, RnlA^18^, ApeA^19^, PrrC, the eukaryotic 2′–5′-oligoadenylate synthetase pathway, and more^17^. Cas13 has been engineered into a versatile platform for programmable RNA knockdown, transcript editing, and nucleic acid diagnostics, demonstrating the substantial biotechnological potential of HEPN-based RNases^20,21^. Yet, how the type IX retron HEPN is inhibited and subsequently activated upon infection, how the WH protein participates in phage defense, and what phage-encoded signal activates the system have not been determined for any type IX retron. Furthermore, all structurally characterized retrons to date have come from *Escherichia coli*.

Here, using cryo-EM, biochemistry, computational protein design, and *in vivo* functional assays, we determined the structure and mechanism of retron-Kva2, a type IX retron from the opportunistic human pathogen *Klebsiella variicola*. We demonstrate that retron-Kva2 assembles into an asymmetric higher-order complex surrounding a HEPN dimer at its core. We further determined that the asymmetric face of retron-Kva2 creates a binding site for a phage trigger, the phage T5 protein D5, which structurally mimics the retron-Kva2 WH. We exploited this recognition principle to design synthetic retron triggers *de novo* that stimulate retron-mediated growth arrest in living cells. Our findings establish synthetic retron triggers as a new platform for selective bacterial elimination and define a general strategy for combating bacterial pathogens by turning their own defense systems against them.

## Results

### Retron-Kva2 forms a higher-order complex surrounding a HEPN dimer

To characterize the type IX retron defense system from *K. variicola* (retron-Kva2), we cloned the entire operon encoding the HEPN, WH, ncRNA, and RT under both T7 and native promoters (Fig. 1a). We induced expression in *E. coli*, and purified the complex by affinity and size-exclusion chromatography. SDS–PAGE confirmed co-purification of the three protein components at the expected molecular weights of approximately 36 kDa (RT), 27 kDa (WH) and 19 kDa (HEPN) (Fig. 1b). We then interrogated the nucleic acid content of our complex with Proteinase K, RNase A, DNase I and Benzonase and revealed the presence of the product of reverse transcription, the msdDNA (55 nt), and the processed msrRNA (37 nt) (Fig. 1c). Together, these data indicate that retron-Kva2 purifies as a functional ribonucleoprotein complex containing all four operonic components.

**Fig. 1:**
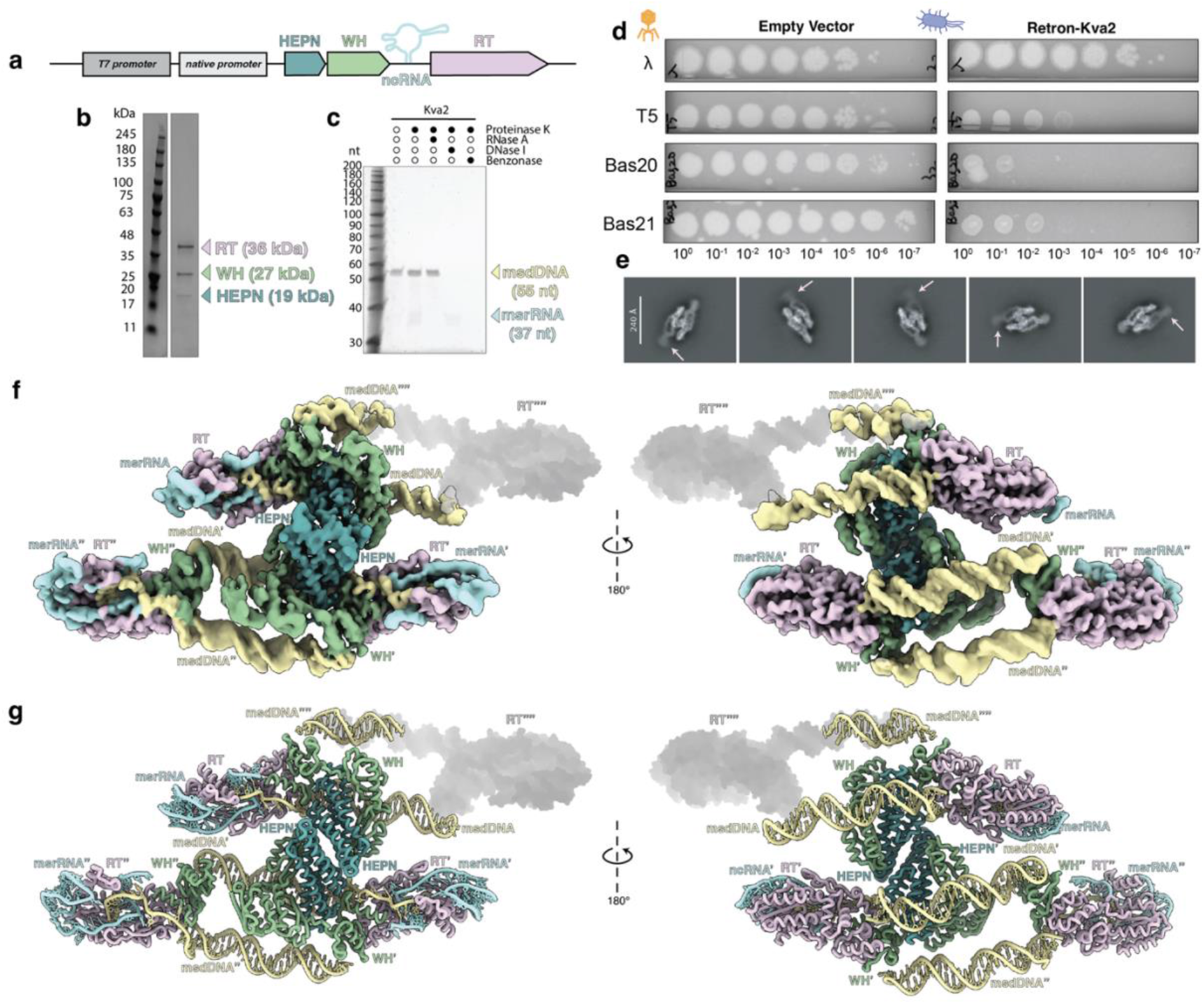
Retron-Kva2 defends against phage and assembles into a higher-order complex around a central HEPN dimer. **(a)** Architecture of the retron-Kva2 operon encoding HEPN, WH, RT and ncRNA under dual T7/native promoter control. **(b)** SDS–PAGE of the purified retron-Kva2 complex showing three co-purifying polypeptides at the expected molecular weights. **(c)** Nuclease-sensitivity analysis identifies the msdDNA (upper band) and processed msrRNA (lower band) as co-purifying nucleic acid species. **(d)** Plaque assays of phages λ, T5, Bas20 and Bas21 on *E. coli* carrying empty vector or retron-Kva2. **(e)** Representative 2D class averages of retron-Kva2; arrows point to a flexible, fourth RT, msrRNA, and msdDNA visible in 2D but averaged out in the high-threshold 3D reconstruction. **(f)** Front and back views of the composite cryo-EM map of the retron-Kva2 complex at a global resolution of 3.7 Å with the fourth RT and msrRNA depicted as a gray shadow. **(g)** Corresponding views of the atomic model built into the composite map.

We next asked whether retron-Kva2 confers phage defense. In plaque assays against a panel of bacteriophage, retron-Kva2 robustly restricted T5, Bas21 and Bas20, but had no detectable effect on phage λ, P1, T7, T4, Bas46, Bas37, Bas15, or Bas1 (Fig. 1d and Extended Data Fig. 1). Intriguingly, T5, Bas20 and Bas21 are all members of the *Markadamsvirinae* subfamily (T5-like phages) that shares a conserved two-step DNA injection mechanism in which ∼8% of the genome initially enters the cell^22^. Retron-Kva2, therefore, restricts a phylogenetically and mechanistically coherent group of phages. Taken together, retron-Kva2 functions as an anti-phage defense system with measurable specificity for T5-like phage with two-step DNA injection mechanisms.

To identify the active composition of retron-Kva2, we determined its structure by cryo-electron microscopy (cryo-EM). 2D class averages readily revealed a higher-order complex that is larger than any one component of retron-Kva2 on its own (Fig. 1e and Extended Data Fig. 2). Subsequent 3D reconstruction, composite map generation, and atomic model building illustrated that retron-Kva2 assembles around a central HEPN homodimer, with three RTs and corresponding ncRNAs, three WHs, and three msDNAs resolvable at a global resolution of 3.7 Å (Fig. 1f-g, Extended Data Fig. 2, and Extended Data Table 1). In the 2D class averages, a fourth RT, msrRNA and msdDNA are visible (arrows, Fig. 1e), but were averaged out in the high-resolution reconstruction (shadows, Fig 1f-g), indicating that the fourth RT and its bound msrRNA and msdDNA are present but conformationally flexible. Local refinement of the flexible fourth RT region did not resolve a fourth WH, reflecting the conformational asymmetry of the complex (Extended Data Fig. 3). The overall stoichiometry of 4 RT: 3 WH: 2 HEPN: 4 msrRNA: 4 msdDNA defines an asymmetric, higher-order architecture in which the HEPN effector is sequestered at the center of the complex.

### The conserved HEPN active site and msDNA stem loop are essential for defense

To pinpoint functional regions within the retron-Kva2 complex, we mapped sequence conservation across type IX retrons from diverse bacterial species onto our atomic model (Fig. 2a and Extended Data Fig. 4). Conserved surfaces clustered at the HEPN dimer interface, the msdDNA stem loop, and the WH pocket that cradles the msdDNA stem loop. The conserved msdDNA positions G29 and A28 contact WH residues D197, R200, and K201 (Fig. 2b and Extended Data Fig. 5). In addition, msdDNA positions G26 and A27 are flipped out and interact with WH residues D190 and N193, creating a structure-specific recognition interface between the tip of the msdDNA stem loop and the WH (Extended Data Fig. 5).

**Fig. 2:**
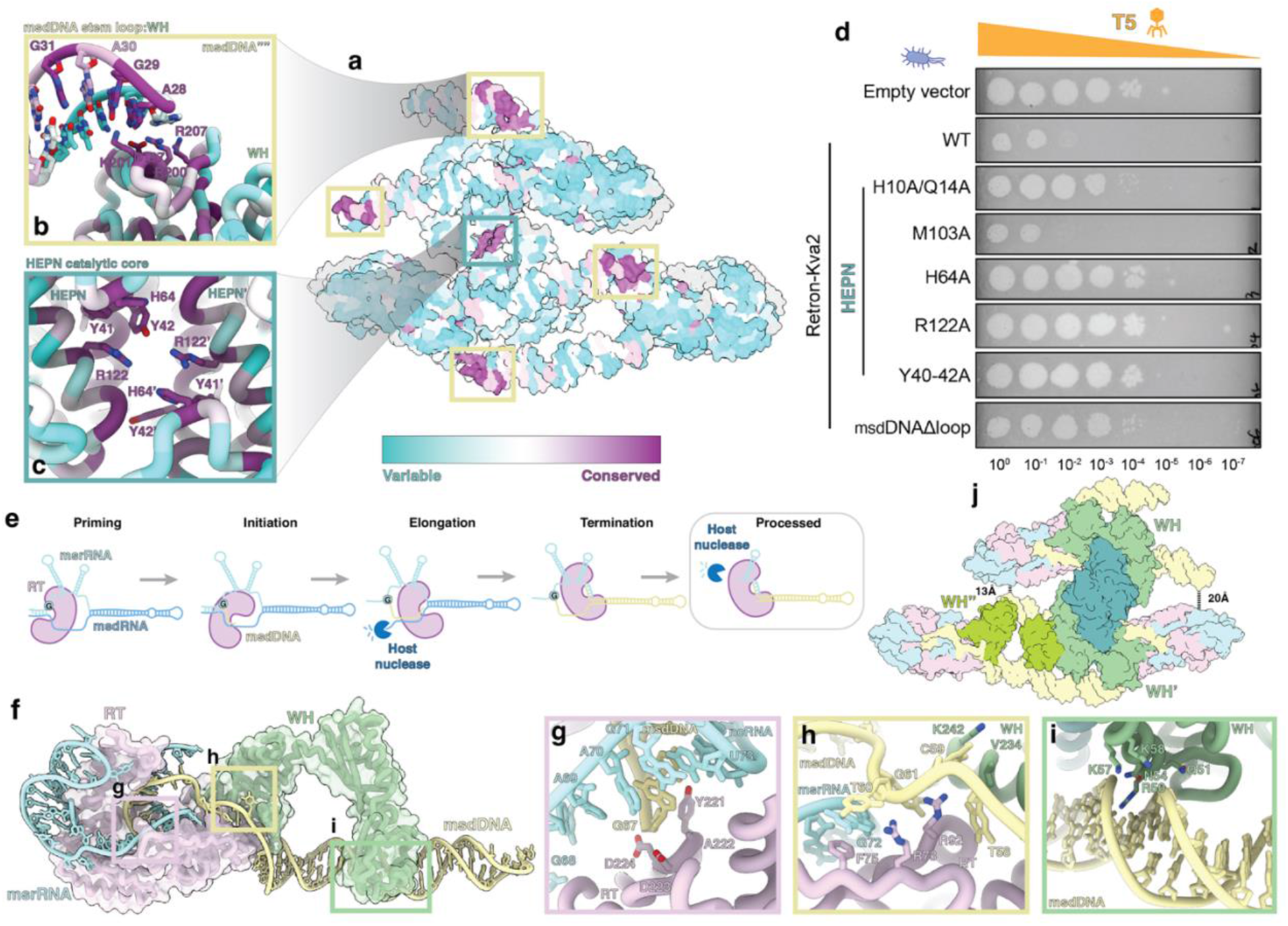
The HEPN catalytic core and the msDNA stem loop are essential for retron-Kva2 phage defense. **(a)** Sequence conservation mapped onto the retron-Kva2 model identifies the HEPN dimer interface and the WH-msdDNA stem loop pocket as the most conserved regions. **(b)** Close-up of the WH-msdDNA interface; nucleotides G29 and A28 contact WH residues R200 and K201. **(c)** Close-up of the HEPN dimer active site showing the symmetric arrangement of catalytic residues H64, R122, H64′, R122′ with substrate-positioning residues Y41 and Y42. **(d)** Plaque assay of T5 on *E. coli* expressing empty vector, WT retron-Kva2, or the indicated mutants. **(e)** Retron msDNA biogenesis pathway. **(f)** A single RT–WH–msDNA protomer of the complex, excluding the central HEPN dimer. **(g)** Close-up of the RT active site cradling the msDNA product with catalytic residues Y221-A222-D223-D224. **(h)** Close-up of the WH alpha helix-msdDNA contacts proximal to the RT. **(i)** Close-up of the WH-msdDNA contacts along the msdDNA stem distal to the RT. **(j)** Structural asymmetry reduces WH occupancy; 13 Å spacing on the two-WH face vs. 20 Å spacing on the single-WH face.

While many HEPN domains are defined by the canonical RϕXXXH, where ϕ is any hydrophobic residue and X is any residue, we could not locate any such sequence in the retron-Kva2 HEPN polypeptide. Comparison across other type IX retron HEPN domains yielded no canonical HEPN motif (Extended Data Fig. 4). Our conservation analysis pointed us toward the HEPN dimer interface, where sequentially distant H64 and R122 on each HEPN monomer (H64, R122, H64′, R122′) spatially converge to form a single active site (Fig. 2c and Extended Data Fig. 5). Together, these residues create a symmetric catalytic core, with Y40-42 from each monomer positioned to engage RNA substrate. Despite the sequential divergence, this geometry is characteristic of HEPN RNases and suggest that retron-Kva2 protects via HEPN-based RNA cleavage.

To functionally validate our structural findings, we generated a panel of retron-Kva2 mutants and tested T5 defense with plaque assays (Fig. 2d). Consistent with our previous results, wild-type retron-Kva2 conferred robust defense. Mutation of the conserved, putative catalytic residues H64A or R122A abolished defense, as did the substrate-positioning triple mutant Y40-42A, and the deletion of the msdDNA stem loop (msdDNAΔloop). The H10A/Q14A double mutant produced an intermediate defect, suggesting that H10 and Q14 contribute to, but are not strictly required, for effector function. Mutation of the non-conserved residue M103 (M103A) had no measurable effect. Thus, both the HEPN active site at the dimer interface and the msdDNA stem loop are essential for retron-Kva2 defense.

Retron ncRNA is highly structured and serves as a scaffold for cognate RT binding. The retron RT binds to the inverted repeat region and initiates reverse transcription at the priming guanosine (Fig. 2e)^23^. The msd region of the ncRNA is reverse transcribed, and the template msdRNA is degraded by host RNase H, yielding in the msDNA^24,25^. Close inspection of the retron-Kva2 structure revealed that the RT is in a termination state, with the msDNA fully processed (Fig. 2f). The RT active site, comprising the canonical YADD (Y221, A222, D223, D224) group II intron motif, cradles the msdDNA product (Fig. 2g and Extended Data Fig. 5). Adjacent to the terminated RT, the WH alpha helix (residues 230-245) nestles against the msdDNA product at positions T57, T58, and C59 (Fig. 2h and Extended Data Fig. 5). The msdDNA residue T60 base-stacks with the msrRNA residue G72 and RT F75, marking the boundary between the msdDNA base-paired with the msrRNA within the RT core and the stem loop region of the msdDNA (Fig. 2h and Extended Data Fig. 5). Along the msdDNA stem, WH residue R50 wedges in the minor grove, while residues Q51, N54 and K57 contact the msDNA backbone (Fig. 2i and Extended Data Fig. 5). Hence, the WH forms a protein bridge between the msdDNA product and the RT active site.

Strikingly, the higher-order complex features an asymmetric distribution of WH proteins, where two WH copies occupy the lower face of the complex (WH’ and WH”) and only one WH copy is bound to the top face (WH)(Fig. 2j). The difference in distance between msdDNA’ C26 and msrRNA U65 provides the structural basis of asymmetry. msdDNA’ and msrRNA are 13 Å from each other, whereas msdDNA and msrRNA’ are 20 Å apart. The 7 Å difference prevents the fourth WH copy from stably binding. Lack of the fourth WH results in increased RT”” flexibility, creating a structurally distinct site that we reasoned could be the location of phage trigger binding.

### Phage T5 protein D5 triggers retron-Kva2-mediated abortive infection

To identify the phage-encoded activator of retron-Kva2, we first expressed retron-Kva2 under its endogenous promoter in an *E. coli* strain lacking its native retron. T5 phages were spotted, and individual plaques were picked and propagated against the cells expressing retron-Kva2. Two independent escapers, T5.1 and T5.2, gained the ability to form plaques in the presence of retron-Kva2 (Fig. 3a), indicating loss-of-function mutations in the phage gene(s) recognized by the retron. Nanopore sequencing of T5.1 and T5.2 surfaced mutations in phage DNA binding, helicase, polymerase, and terminase genes (Extended Data Table 2 and Extended Data Fig. 6). Co-cultures of plasmids expressing these phage genes with retron-Kva2, or empty vector indicated phage genes D2, D3, and D5 affected cell growth in the presence of retron-Kva2 (Extended Data Fig. 6). T5.1 carried four mutations in D5, G53D, a premature stop codon W74*, V149I, and G232D, as well as an E76Q substitution in D2 (Fig. 3b). T5.2 carried E30Q and G85D mutations in D3 (Fig. 3b). We next tested whether T5 D2, D3, or D5 could trigger retron-Kva2-mediated defense in the absence of phage infection by simply co-expressing either an empty vector or retron-Kva2 (expressed from its native promoter) together with a candidate phage trigger, with the trigger placed under an arabinose-inducible promoter (Fig. 3c). T5 D2 exhibited inherent toxicity, regardless of the presence of retron-Kva2. In contrast, T5 D3 did not induce growth arrest in the presence of retron-Kva2, although we did observe a mild effect on colony morphology. T5 D5 displayed toxicity on its own, but the toxicity was amplified in cells expressing retron-Kva2, suggesting that T5 D5 is a trigger of retron-Kva2 defense.

**Fig. 3:**
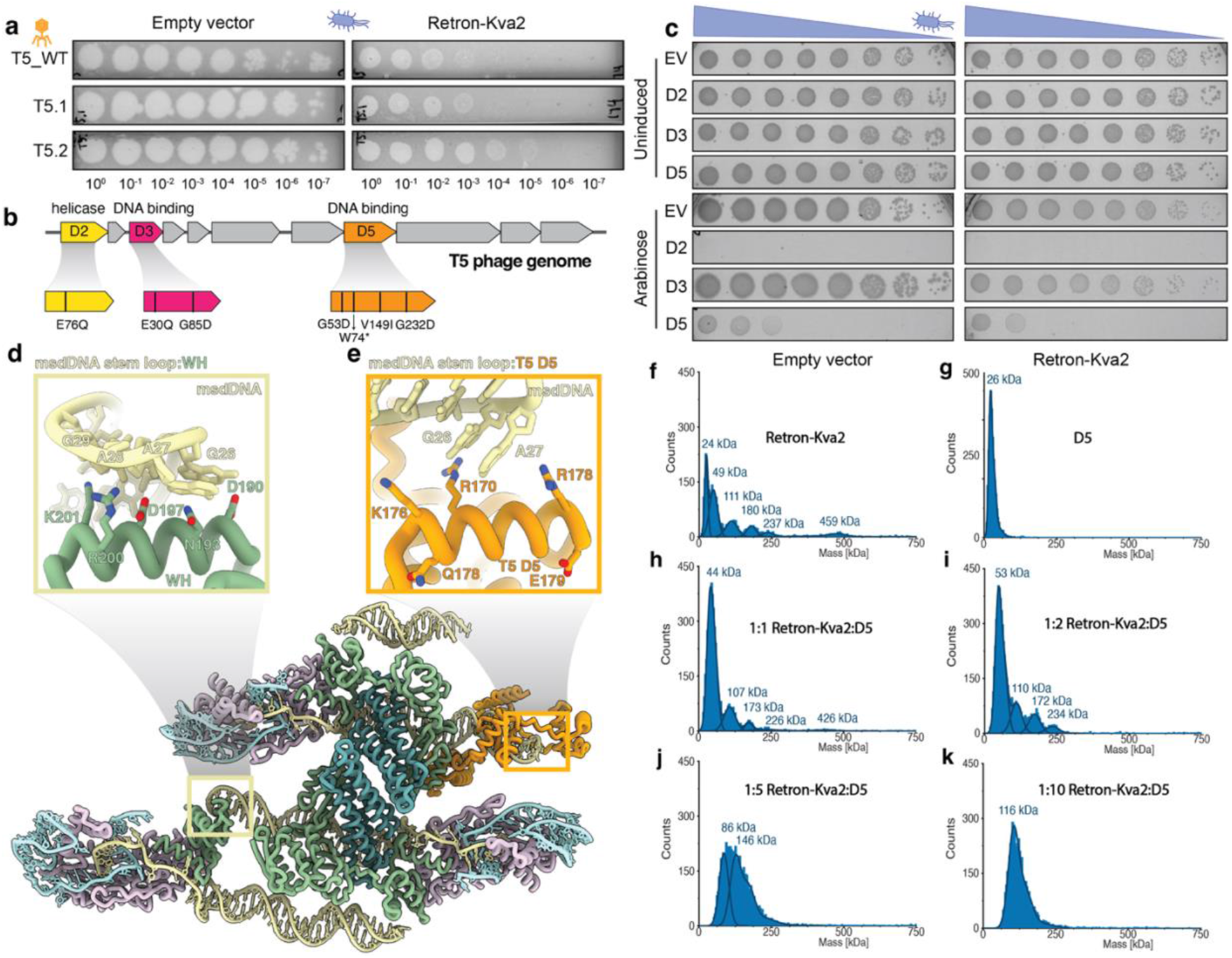
Phage T5 protein D5 triggers retron-Kva2 by molecular mimicry of the WH domain. **(a)** Plaque assays showing T5 escape mutants T5.1 and T5.2 that regain plaquing on retron-Kva2-expressing *E. coli*. **(b)** Mutational mapping of T5.1 and T5.2 escape phages identifies mutations in the DNA binding (D5, D3) and helicase (D2) genes. **(c)** Co-expression spot dilution assays of uninduced and arabinose-induced T5 proteins (D2, D3, D5) with retron-Kva2 or an empty vector. **(d)** Close-up of the experimentally resolved WH-msdDNA stem loop interface. **(e)** Close-up of the AlphaFold3^26^ model of T5 D5 and msdDNA, which was docked onto the experimentally resolved retron-Kva2 complex, highlighting the interaction mimicry. **(f–k)** Mass photometry analysis of Retron-Kva2 alone **(f)**, D5 alone **(g)**, and Retron-Kva2:D5 mixtures at molar ratios of 1:1 **(h)**, 1:2 **(i)**, 1:5 **(j)**, and 1:10 **(k)**, demonstrating partial complex disassembly.

To understand the molecular basis of D5 recognition, we used AlphaFold3^26^ to predict the structure of T5 D5 and retron-Kva2 msdDNA, in conjunction with our other potential phage triggers, as well as other known phage triggers of different retron systems (Extended Data Fig. 7). Only D5 was predicted to bind to retron-Kva2 msdDNA. D5 contains a helix-turn-helix (HTH) motif that engages the msdDNA in a manner highly analogous to the retron-Kva2 WH. Comparison of the WH-msdDNA interface with the predicted D5-msdDNA interface shows a nearly identical amino acid composition, including arginines and lysines that foster electrostatic interactions with the msdDNA phosphate backbone, as well as negatively charged aspartate and serine residues that impart specificity for the stem loop region of the msdDNA (Fig. 3d-e). Structural alignment of D5 and the retron-Kva2 WH highlights their structural similarity, with an RMSD of 1.3 Å (Extended Data Fig. 7). We propose that D5 activates retron-Kva2 by molecular mimicry, where D5 binds to the asymmetric face of the retron complex and triggers HEPN activation.

To further characterize D5 engagement with retron-Kva2, we performed mass photometry experiments where we incubated both with varying stoichiometric ratios, as well as D5 and retron-Kva2 on their own (Fig. 3f). Retron-Kva2 alone exhibited multiple size species, ranging from approximately 24 kDa to 459 kDa. Each species corresponds to a complex, subcomplex, or subunit of retron-Kva2, with the largest peak corresponding to the higher-order architecture we resolved by cryo-EM.

On its own, D5 was seen at its expected size of 26 kDa (Fig. 3g). Interestingly, combining D5 with retron-Kva2 depleted the higher-order species in a concentration-dependent manner, with an appreciable difference when D5 and retron-Kva2 were mixed at equimolar concentrations (Fig. 3h). When we doubled the concentration of D5 relative to retron-Kva2, we were no longer able to detect the largest higher-order species, with the signal redistributing toward an intermediate-sized population (Fig. 3i). A stark transition emerged when we increased the concentration of D5 to five times that of retron-Kva2, and one broad peak spanning 80-150 kDa appeared (Fig. 3j). At a ratio of 10:1, the population centered at a peak of around 116 kDa (Fig. 3k). The concentration-dependent disappearance of the higher-order architecture indicates that D5 does not simply dock onto the complex, but actively remodels it. Of note, this remodeling is not a total disassembly, as has been described for other retron systems^27^, but rather, D5 binding yields a partial, likely conformationally rearranged ∼116 kDa subcomplex that we propose represents the catalytically active state of retron-Kva2.

### Computational design of synthetic retron triggers

The structural mimicry between phage T5 protein D5 and the host WH protein suggested that a protein capable of binding to the msdDNA stem loop should, in principle, activate retron-Kva2. To test this hypothesis directly, we developed a computational pipeline to design *de novo* synthetic trigger proteins (Fig. 4a). The pipeline proceeded in six steps. First, the AlphaFold3-predicted^26^ structure of T5 D5 was used as the seed query. Second, a FoldSeek^28^ search against the AlphaFold Database^29^ revealed HTH-containing proteins spanning from bacterial D5 homologs, to eukaryotic transposases and transcription factors (Extended Data Fig. 8). The phylogenetic breadth of these hits frames the HTH motif as a conserved scaffold for nucleic acid recognition, suggesting that direct msdDNA binding is the primary trigger for type IX retron activation. Third, multiple sequence alignment^30^ of the hits yielded a consensus sequence capturing the minimal HTH-fold features (Fig. 4b). Fourth, RFdiffusion^31^ was used in partial-diffusion mode in which D5 residues at the predicted msdDNA-binding interface were retained as fixed contigs while connecting and flanking segments were diversified (Fig. 4b-c). These partial-diffusion steps produced *de novo* backbones that preserved the HTH geometry and the D5-derived binding surface. Fifth, ProteinMPNN^32^ was used to design amino acid sequences optimized for stability and solubility on each generated backbone (Fig. 4d). Sixth, AlphaFold3^26^ was used to predict the structures of each designed trigger in complex with the retron-Kva2 msdDNA (Fig. 4e and Extended Data Fig. 9). This pipeline yielded six synthetic trigger candidates, designated 7n7, 7n5, 5n6, 7n6, 7n3 and 2n0. All six designs present a positively charged surface engaged with the msdDNA stem loop that is akin to the electrostatic surface potential of D5 (Fig. 4f). By distilling the functional interface into a 126-amino-acid scaffold, which is roughly half the 255-residue footprint of natural D5, these *de novo* designs offer a compacted architecture that is highly advantageous for translation into size-constrained therapeutic delivery systems. The shared binding modes of our synthetic triggers with D5 suggested that our designs could trigger retron-Kva2 *in vivo*.

**Fig. 4:**
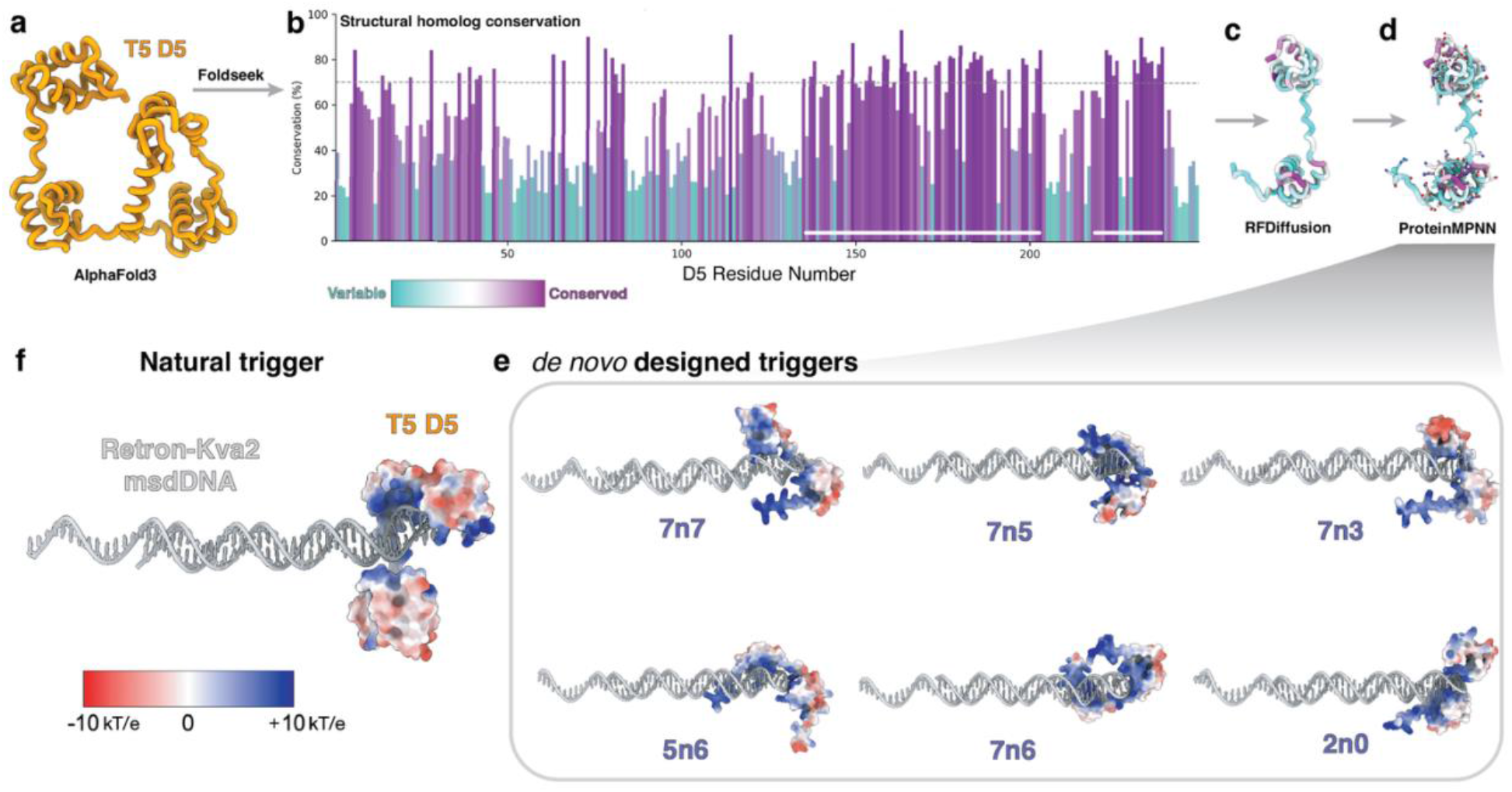
Computational pipeline used to generate synthetic retron triggers. **(a)** AlphaFold3^26^ structural model of T5 D5. **(b)** Sequence conservation of D5 structural homologs, mapped onto the D5 sequence; the msdDNA interface segments (contigs) retained for RFdiffusion^31^ indicated with white bars. **(c, d)** Computational design steps utilizing RFDiffusion for partial-diffusion backbone generation **(c)** followed by ProteinMPNN^32^ for sequence design **(d)**. **(e)** Electrostatic surface representations of six *de novo* designed synthetic triggers (7n7, 7n5, 7n3, 5n6, 7n6, 2n0) predicted to bind to retron-Kva2 msdDNA by AlphaFold3. **(f)** Electrostatic surface representation of the T5 D5 predicted to bind to retron-Kva2 msdDNA by AlphaFold3.

### Synthetic retron triggers activate retron-Kva2 in vivo

To evaluate our computational designs *in vivo*, we co-expressed the T5 D5 trigger or our synthetic variants (5n6 or 7n7) with either retron-Kva2 or an empty vector in *E. coli* (Fig. 5a). Induction of the D5 protein alone caused mild inherent toxicity, consistent with our previous results, but co - expression with retron-Kva2 amplified this effect to drive robust cellular elimination (Fig. 5b). Crucially, both *de novo* designs were non-toxic in the absence of retron-Kva2, yet they stimulated potent and targeted growth arrest when co-expressed with retron-Kva2. The 7n7 design proved exceptionally sensitive, reducing cell viability even under uninduced conditions due to basal leaky expression, and eliciting similar growth arrest upon full induction. Although D5 provoked the most substantial decrease in cell viability, both synthetic designs significantly compromised cell survival upon induction, with 7n7 demonstrating greater potency than 5n6. These results establish that retron-Kva2 activation is governed by the structural presentation of the msdDNA-binding surface, and highlights the advantage of computationally derived triggers in eliminating off-target host toxicity.

**Fig. 5:**
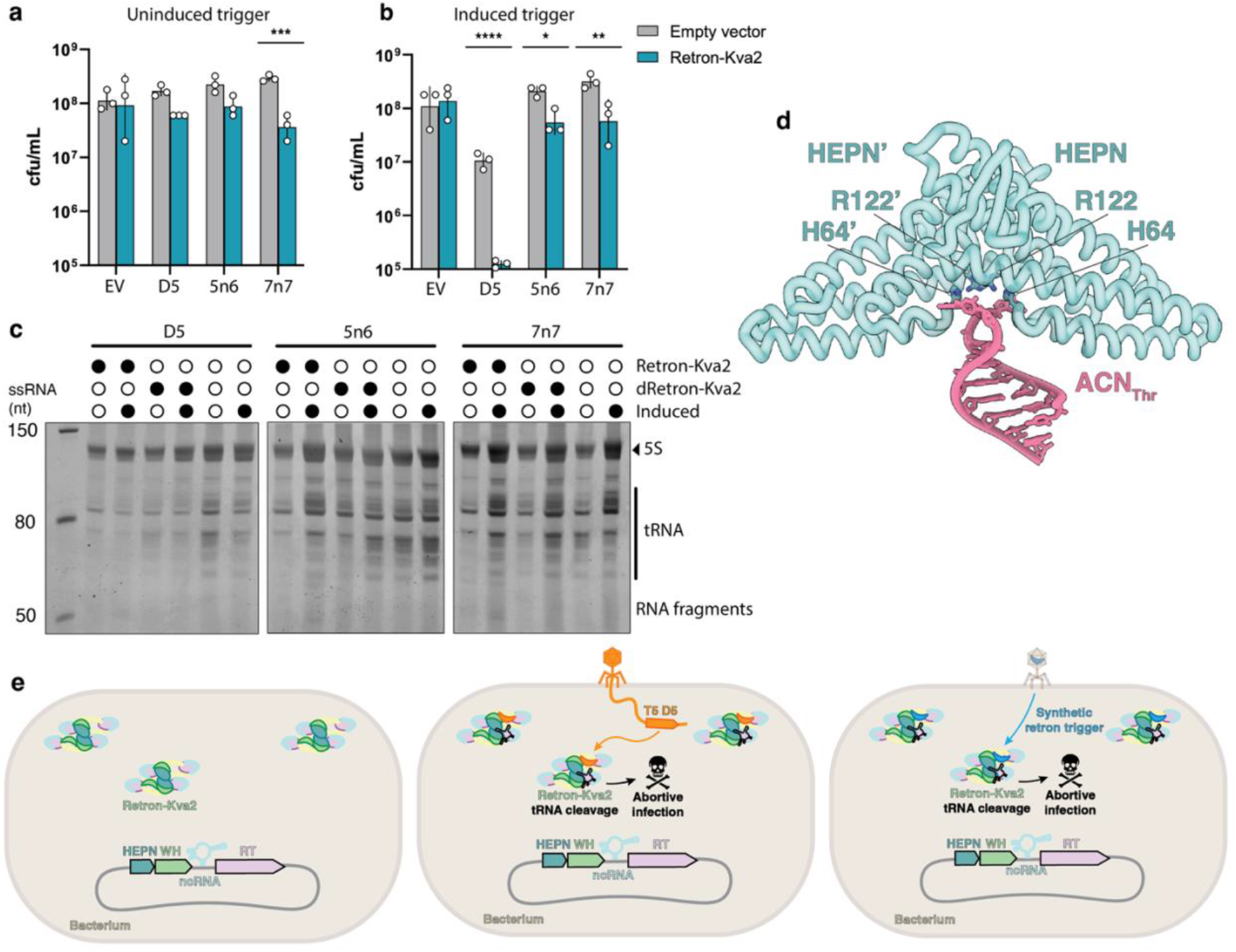
Natural and synthetic triggers activate retron-Kva2 HEPN tRNase activity *in vivo*. Bacterial cell viability assay testing the effects of D5 and synthetic triggers 5n6 and 7n7 under **(a)** uninduced trigger, and **(b)** induced trigger conditions. *n* = 3 biological triplicates. Significance was determined using a two-way ANOVA. **(c)** Total RNA extracted from *E. Coli* cultures co-expressing retron-Kva2, catalytically dead retron-Kva2 (HEPN H64A/R122A), or empty vector, and D5, 5n6, or 7n7 three hours post-induction. **(d)** AlphaFold3 model of the HEPN dimer active site bound to the tRNA^Thr^ anticodon stem loop (ACN_Thr_) **(e)** Proposed mechanism of retron-Kva2 defense: under uninfected conditions, retron-Kva2 is constitutively expressed forming higher-order asymmetric complexes. Upon phage infection, D5 binds to the asymmetric binding site leading to partial complex disassembly, HEPN activation, tRNA degradation, and abortive infection. We propose that our synthetic triggers act similarly on retron-Kva2, stimulating cell growth arrest.

We next investigated the downstream mechanism of retron-Kva2-mediated abortive infection. Total RNA profiling three hours post-induction revealed that D5 expression triggered a substantial depletion of tRNA-sized species in the presence of retron-Kva2, and even in the presence of catalytically dead retron-Kva2 (HEPN H64A/R122A) albeit to a lesser degree, which is likely due to the innate toxicity of D5 we have observed across all our experiments (Fig. 5c). Chemical induction increases the quantity of tRNA in the cell, as seen in our RNA extractions from 5n6 and 7n7 co-cultures. Despite more total tRNA under induced conditions, smaller RNA fragments appear only when the triggers were induced with retron-Kva2, and not with catalytically dead retron-Kva2. Without retron-Kva2, tRNA remain intact in the presence of 5n6 or 7n7, although D5 stimulates tRNA depletion on its own when induced. Together, these data not only indicate that retron-Kva2 executes abortive infection through tRNA degradation, but also that retron-Kva2-mediated abortive infection can be activated by *de novo* computationally designed retron triggers.

To identify plausible RNA substrates, we looked to the established biology of HEPN RNases. In diverse bacterial defense systems including CRISPR-Cas13, HEPN RNases frequently execute cell dormancy by specifically targeting and cleaving tRNA anticodon stem loops to halt host translation^33–35^. Guided by well-characterized HEPN targets, which exhibit preferences for uridine-rich anticodons^35^, we leveraged AlphaFold3^26^ to model these specific candidate anticodon stem loops with the retron-Kva2 HEPN dimer. While AlphaFold3 predicted binding for both a lysine tRNA anticodon stem loop (ACN_Lys_) and a phenylalanine tRNA anticodon stem loop (ACN_Phe_), neither was positioned within the HEPN catalytic active site at the dimer interface (Extended Data Fig. 9). In contrast, a threonine tRNA anticodon stem loop (ACN_Thr_) docked in a catalytically competent orientation between the conserved H64 and R122 active site residues from each HEPN monomer (Fig. 5d and Extended Data Fig. 10). When overlaid with our experimentally determined model of the HEPN dimer, the predicted HEPN dimer with ACN_Thr_ reveals a 3 Å shift within the active site (Extended Data Fig. 10). This shift is likely the molecular basis of retron-Kva2 activation, gating collateral tRNA cleavage prior to phage infection. Although the precise tRNA target remains to be experimentally validated, the predicted structural discrimination indicates the retron-Kva2 HEPN may possess anticodon specificity. Integrating these structural and *in vivo* findings, we propose a model in which retron-Kva2 senses trigger binding at its asymmetric face via the msdDNA stem loop, prompting a conformational change in the HEPN dimer that leads to tRNase activity and abortive infection.

## Discussion

Our structural and functional characterization of retron-Kva2 establishes a mechanistic framework for type IX retron phage defense and reveals a programmable recognition principle for engineered cellular elimination. Retron-Kva2 assembles into an asymmetric higher-order ribonucleoprotein complex held in an autoinhibited state, where WH occupancy underlies the architectural asymmetry. The asymmetric WH binding makes one face of retron-Kva2 more compacted than the opposite face, creating an exposed binding site at the msdDNA stem loop. Phage T5 protein D5 docks at this msdDNA interface, which a fourth WH cannot stably bind to due to the wider spacing. D5 binding initiates partial complex disassembly that culminates in HEPN tRNase activity at the core of the complex. Thus, retron-Kva2 is organized so that catalysis is coupled to recognition, with the architectural asymmetry positioning the HEPN dimer for prompt activation upon direct msdDNA engagement.

We identified phage T5 protein D5 as a trigger of retron-Kva2. Our functional assays and *de novo* designs demonstrate that activation relies on the topological recognition of the msdDNA stem loop. By utilizing the msdDNA as a sensory scaffold, retron-Kva2 acts as a direct molecular sensor for specific protein structures. Specificity derives from physical docking at the msdDNA, which prompts the conformational change required for HEPN activation.

This unique protein-sensing capability provides a compelling platform for the rapidly expanding field of programmed cell death. Recent landmark studies have demonstrated the therapeutic potential of repurposing bacterial abortive infection systems, notably through reprogramming the RNA-guided nuclease Cas12a2 to sense mutant transcripts and drive selective cell death in precancerous models^36,37^. While these foundational Cas-mediated technologies highlight the clinical promise of harnessing host cell death, they fundamentally rely on nucleic acid recognition. Retrons offer a transformative departure by directly detecting proteins.

The vast phylogenetic diversity of retrons^12,38^ represents a naturally generated arsenal of defense systems already tuned to sense a myriad of distinct phage proteins^27,39–41^. Recent structural studies of retron complexes highlight this evolutionary divergence with msdDNA topologies ranging from stem loops to kinked single-stranded DNA architectures^23,27,39,40,42–46^. The structural plasticity of the msdDNA is likely the primary determinant for distinguishing diverse phage proteins. By rationally re-engineering the structural folds of the msdDNA, we could theoretically tune retrons to bind and sense user-defined protein targets that lack trackable nucleic acid signatures.

As proof of concept, we show that synthetic polypeptides generated via a structure-guided computational pipeline activate retron-Kva2-mediated growth arrest in *E. coli*. Unlike the phage D5 protein, which exhibits inherent toxicity to the host, our *de novo* synthetic triggers are non-toxic when expressed in isolation, thereby significantly widening the therapeutic window for engineered applications. Furthermore, our designs are half the size of the D5. This compact size is highly advantageous for size-constrained delivery systems such as viral vectors, where cargo capacity is often the limiting factor. By defining the molecular interface sensed by retron-Kva2, and recapitulating it with programmable synthetic triggers, we transition from describing an immune mechanism to actively programming it. The biochemical diversity of bacterial defense systems serves not just as a catalogue of evolutionary interest, but as a highly modular toolkit from which entirely new classes of protein-responsive therapeutics can be built.

## Author contributions

G.N.H. conceptualized the project and designed the study. G.N.H. and L.W. developed the methodology. L.W. constructed all plasmids used for the study. G.N.H. and N.M. generated mutant plasmids. G.N.H. performed protein production, cryo-EM data collection, data analysis, atomic modeling, computational design of synthetic triggers, AlphaFold predictions and analysis, and total RNA profiling. L.W. performed phage defense assays, phage escapee isolation and analysis, trigger co-expression assays, and growth curves. N.M. performed nucleic acid composition assays, conservation analyses, and mass photometry. K.Z. performed and analyzed trigger co-expression assays and assisted with phage assays. A.F. assisted with cryo-EM data analysis strategy and atomic modeling. G.N.H. wrote the original draft of the manuscript. G.N.H., L.W., N.M., and K.Z. created the data visualizations. All authors reviewed, edited, and approved the manuscript. G.N.H., S.L.S., and E.N. acquired funding and supervised the research.

## Methods

### Bacterial Strains and Phages

Routine cloning was performed in *E. coli* strain DH5α. Protein expression was conducted in *E. coli* BL21 (DE3). Phage defense and trigger validation experiments were conducted in a derivative of *E. coli* BL21-AI lacking its native Retron-Eco1 (bSLS.114)^47^. Bacteriophages (P1_vir_, T5, λ_vir_, T7, T4, Bas51, Bas46, Bas37, Bas21, Bas20, Bas15, and Bas1) were propagated in *E. coli* bSLS.114 liquid cultures at 37°C. Cultures were grown in MMB media (LB supplemented with 0.1 mM MnCl_2_ and 5 mM MgCl_2_). Following overnight propagation, phage lysates were isolated by centrifugation (10 min at 3,434*g*) and syringe filtration of the supernatant through a 0.2-micron filter.

### Plasmid Construction and Mutagenesis

The wild-type *retron-Kva2* operon (encoding the HEPN, WH, RT, and ncRNA) from *Klebsiella variicola* under the control of dual T7 and native promoters was synthesized as gene block from Twist Bioscience and was assembled into digested pMRM032, generating pLW174. Point mutations in retron-Kva2 HEPN (H64A, R122A, H10A, Q14A, M103A, Y40-42A), and the msdDNA loop deletion (msdDNAΔloop) were generated using Q5 Hot Start High-Fidelity mutagenesis (New England Biolabs). Phage trigger genes (pLW183-189) were amplified from the isolated T5 phage genome and were assembled into the digested pKAZ208. All the phage triggers are listed in the Extended Data Table 2. All constructs were verified by whole plasmid sequencing using Oxford Nanopore Technology performed by Plasmidsaurus.

### Protein Expression and Purification

*E. coli* cells harboring the retron-Kva2 expression plasmid were grown to an optical density OD_600_ = 0.8 and induced with 0.2 mM IPTG for 18 hours at 18°C. Cells were harvested by centrifugation, resuspended in lysis buffer (20 mM HEPES pH 7.5, 300 mM NaCl, 5% glycerol, 5 mM MgCl₂, 1 mM TCEP), supplemented with protease inhibitors, and lysed via sonication. The lysate was cleared by centrifugation, and the retron-Kva2 complex was isolated using Ni-NTA affinity chromatography. Following washing with a high salt wash buffer (20 mM HEPES, 1 M NaCl, 20 mM imidazole, 5 mM MgCl₂, 5 mM ATP, 5% glycerol, 1 mM TCEP), the complex was eluted using buffer containing (20 mM HEPES pH 7.5, 300 mM NaCl, 250 mM imidazole, 5 mM MgCl₂, 5% glycerol, 1 mM TCEP). The eluate was buffer exchanged through a 30 kDa spin concentrator into size-exclusion chromatography (SEC) buffer (20 mM HEPES pH 7.5, 150 mM NaCl, 5% glycerol, 5 mM MgCl₂, 1 mM TCEP) and further purified by SEC using a Superdex 200 Increase 10/300 GL column equilibrated in SEC buffer. Protein concentrations were estimated using the Warburg-Christian method, calculated via the equation C = 1.55 x A_280_ - 0.76 x A_260_, to account for nucleic acid absorbance within the ribonucleoprotein complex. Co-purification of the HEPN, WH, and RT components was confirmed via SDS–PAGE. The phage trigger D5 was purified utilizing an analogous Ni-NTA and SEC pipeline.

### Nucleic Acid Composition Assays

To determine the nucleic acid composition of the purified retron-Kva2 complex, 300 ng of the purified complex were treated with either 1 μL of Proteinase K (1 mg/mL), RNase A (1 mg/mL), DNase I (2 U/μL), or Benzonase (25 U/μL) at 37°C for 20 minutes. Reactions were quenched with 0.5 M EDTA, dyed with 2X TBE-Urea buffer (Invitrogen), and boiled for 10 minutes at 90°C. The resulting species were resolved on a 10% Urea-PAGE gel stained with SYBR Gold (Pierce Thermo Scientific) and visualized under UV transillumination to identify the 55 nt msdDNA and 37 nt msrRNA.

### Phage Defense Assays

For phage defense assays, *E. coli* bSLS.114 cells expressing either an empty vector or wild-type/mutant retron-Kva2 under its native promoter were grown to exponential phase in MMB media. 200 µL of this culture was then mixed with 3 mL MMB top agar (0.75% agar in MMB) and poured onto a MMB agar plate (containing 1.5% agar in MMB). After the top agar solidified, ten-fold serial dilutions of phage lysates were spotted onto the bacterial lawns in 2 µL volumes and incubated overnight at 37°C.

### Phage Escaper Isolation and Sequencing

Isolated T5 plaques that formed on retron-Kva2-expressing lawns in previous spot assays were picked and resuspended in 90 µL SM buffer (200 mM NaCl_2_, 10 mM MgSO_4_, 50 mM Tris-HCl pH 7.5). This resuspension was briefly vortexed to disperse the agar and left at 4 C for at least 4 hours to allow phage to diffuse into the buffer. The entire resuspension was then used to infect a culture of retron-Kva2-expressing cells at an OD600 of 0.3 and grown overnight. Lysates were isolated and then spotted onto a lawn of the retron-expressing bacteria (or empty vector bacteria as a control).

For the T5 escapers that successfully plaqued on the retron-expressing bacteria, we performed whole genome sequencing. Genomic DNA from wild-type T5 and escaper mutants (T5.1, T5.2) was extracted using Norgen’s Phage DNA Isolation Kit (Norgen Biotek 46800). Purified genomic DNA was prepped for sequencing using Oxford Nanopore Technologies’ Native Barcoding Kit (SQK-NBD114.96) and sequenced on a PromethION P2I machine using a R10.4.1 flow cell (FLO-PRO114M). Reads were aligned to the reference T5 genome to identify loss-of-function single nucleotide polymorphisms and premature stop codons in genes *D2*, *D3*, and *D5*.

### In Vivo Toxicity and Trigger Co-Expression Assays

*E. coli* bSLS.114 cells were co-transformed with retron-Kva2 plasmid and an arabinose-inducible plasmid encoding either D2, D3, D5, synthetic triggers (5n6, 7n7), or an empty vector. Overnight cultures were inoculated into fresh LB for 2 hours to OD600 of 0.4. Ten-fold serial dilutions were spotted 5 µl onto agar plates with or without inducing media (2 mg/mL L-arabinose for triggers) alongside requisite antibiotics. Plates were incubated at 37°C overnight and bacterial colonies were imaged and counted.

### Growth Curve Co-Expression Assays

Retron-Kva2 or an empty vector control was co-transformed with a plasmid containing the trigger gene in *E. coli* bSLS.114. Individual colonies were picked and grown until saturation in 1 mL LB with carbenicillin and chloramphenicol at 37°C. Cultures were then diluted 1:100 in 200 µL of fresh LB with carbenicillin and chloramphenicol in a 96-well microtiter plate. The plate was covered with an air-permeable seal and incubated in a microplate reader with shaking at 37°C for 16 hours with OD_600_ measured every 10 min. At the two-hour timepoint, arabinose was added to all cultures. Raw OD_600_ values were adjusted by subtracting the zero-hour measurement for each sample.

### Total RNA Profiling and tRNA Degradation Assay

*E. coli* BL21 DE3 cells were co-transformed with retron-Kva2, catalytically dead retron-Kva2 (HEPN H64A/R122A), or empty vector and the specified triggers (D5, 5n6, or 7n7). The co-transformed cells were allowed to outgrow in SOC media for 1 hour at 37°C at 220 rpm. The outgrowth was expanded into 5 mL LB with ampicillin (100 μg/mL), chloramphenicol (34 µg/mL), and 1% glucose. The expanded cultures were allowed to grow overnight. The next day, the dense overnight cultures were subcultured 1:25 in fresh LB and corresponding antibiotics and 1% glucose. When the subculutres reached and OD_600_ of 0.6, they were pelleted, washed with LB to remove the glucose, resuspended in 5 mL LB with ampicillin and chloramphenicol, and split in half. One half of the culture was induced with 0.2 mM IPTG, and 0.2% L-arabinose 30 minutes after IPTG induction. The other half was grown without inducers. At three hours post-induction, cells were harvested, and total RNA was extracted using Monarch Total RNA Extraction Kit. Total RNA was quantified, and 300 ng per sample were resolved on a 10% TBE-Urea gel, and stained with SYBR Gold to visualize the depletion of tRNA relative to uninduced controls.

### Mass Photometry

Mass photometry was performed using a Refeyn OneMP to assess complex remodeling upon D5 engagement. Standard glass coverslips were cleaned with isopropanol. Retron-Kva2 and T5 D5 were analyzed individually and mixed at specific molar ratios (1:1, 1:2, 1:5, and 1:10 Retron-Kva2:D5) in SEC buffer. Droplets were measured for 60 seconds. Data were processed and molecular weight distributions were plotted using DiscoverMP.

### Cryo-EM Sample Preparation and Data Acquisition

Purified retron-Kva2 complex at 5.5 μM was applied to glow-discharged Quantifoil R1.2/1.3 Cu 300 mesh grids. Grids were blotted for 6 seconds at 100% humidity at 4°C and plunge-frozen in liquid ethane using a FEI Mark IV Vitrobot. Cryo-EM data were collected on a 200 kV Talos Arctica electron microscope equipped with a Gatan K3 direct electron detector. 17,178 movies were recorded with SerialEM 4.2^48^ at a nominal magnification of 36,000 corresponding to a physical pixel size of 1.14 Å/pix. A total dose 50 e⁻/Å² was applied, with a target defocus range of-1 to-3 µm.

### Cryo-EM Data Processing

Motion correction and CTF estimation were performed using cryoSPARC Live^49^. Initial particle picking was done using a blob picker, followed by 2D classification, *ab initio*, heterogeneous refinement, and non-uniform refinement to generate templates for subsequent automated template-based picking in cryoSPARC (v5.0). Extracted particles were subjected to one round of selective 2D classification to remove junk particles and ice artifacts. The resulting clean particle stack was used to generate *ab initio* maps, followed by heterogeneous refinement using the two best *ab initio* maps duplicated as references. The best heterogeneous refinement class was further refined with non-uniform refinement and exported for subsequent processing in RELION (v5.0)^50^.

The exported particle set was re-refined in RELION using 3D refinement with Blush^50^. The resulting map was used to create masks around the top and bottom lobes of the complex. Each were exposed to particle subtraction, re-centering, and 3D back-projection with a low-pass filter of 10 Å. The subtracted particles were then refined with 3D refinement with Blush. The peripheral RT was then isolated from the bottom lobe, and underwent particle subtraction, re-centering, re-boxing to 300-pixel box size. The peripheral RT particles were the back-projected and low-pass filtered to 10 Å. Each isolated component of the complex (top, bottom, or peripheral RT) were then parsed by 3D classification without alignment with a T-value of 500. A major class emerged from each classification, which was finally refined with 3D refinement with Blush. Each locally refined map was merged into a composite map at a global resolution of 3.7 Å, estimated using the gold-standard Fourier shell correlation (FSC) = 0.143 criterion.

### Atomic Model Building and Refinement

Initial models of the HEPN, WH, RT, and msdDNA subunits were generated using AlphaFold3^26^. These models, along the msrRNA from PDB 7XJG^23^ which was re-sequenced to retron-Kva2 msrRNA, were rigidly docked into the 3.7 Å composite cryo-EM map using ChimeraX v1.11^51^. Flexible fitting and manual rebuilding of the sequence into the density were performed in ISOLDE^52^. Real-space refinement was iteratively carried out in Phenix v2.0^53^ with secondary structure restraints applied. Figures were rendered using ChimeraX v1.11^51^.

### Computational Design of Synthetic Triggers

Structural prediction of the phage T5 protein D5 was conducted using AlphaFold3^26^. To identify structural homologs, this D5 model was used as a search query against the AlphaFold Database using FoldSeek^28^. Identified hits were aligned using MAFFT^30^ to extract the conserved HTH regions of D5. For the generation of *de novo* triggers, we employed RFdiffusion^31^ in partial-diffusion mode. The conserved HTH D5 residues were designated as fixed contigs, while the interconnecting and flanking segments were allowed to diversify. Generated backbones preserving the HTH geometry were subjected to sequence design using ProteinMPNN^32^, optimized for solubility and stability. The resulting sequences (7n7, 7n5, 5n6, 7n6, 7n3, 2n0) were tested by co-folding with retron-Kva2 msdDNA via AlphaFold3.

### Data Availability

The retron-Kva2 composite and consensus structures along with associated focused refinement maps for the bottom lobe, peripheral RT, and top lobe have been deposited to the PDB and EMDB with the accession codes PDB 36HT, EMD-77585, EMD-77583, EMD-77573, EMD-77574, and EMD-77575, respectively.

