## Supplemental Figures and Tables for "Higher-order assembly of a type IX retron enables exploitation for designer antimicrobials"

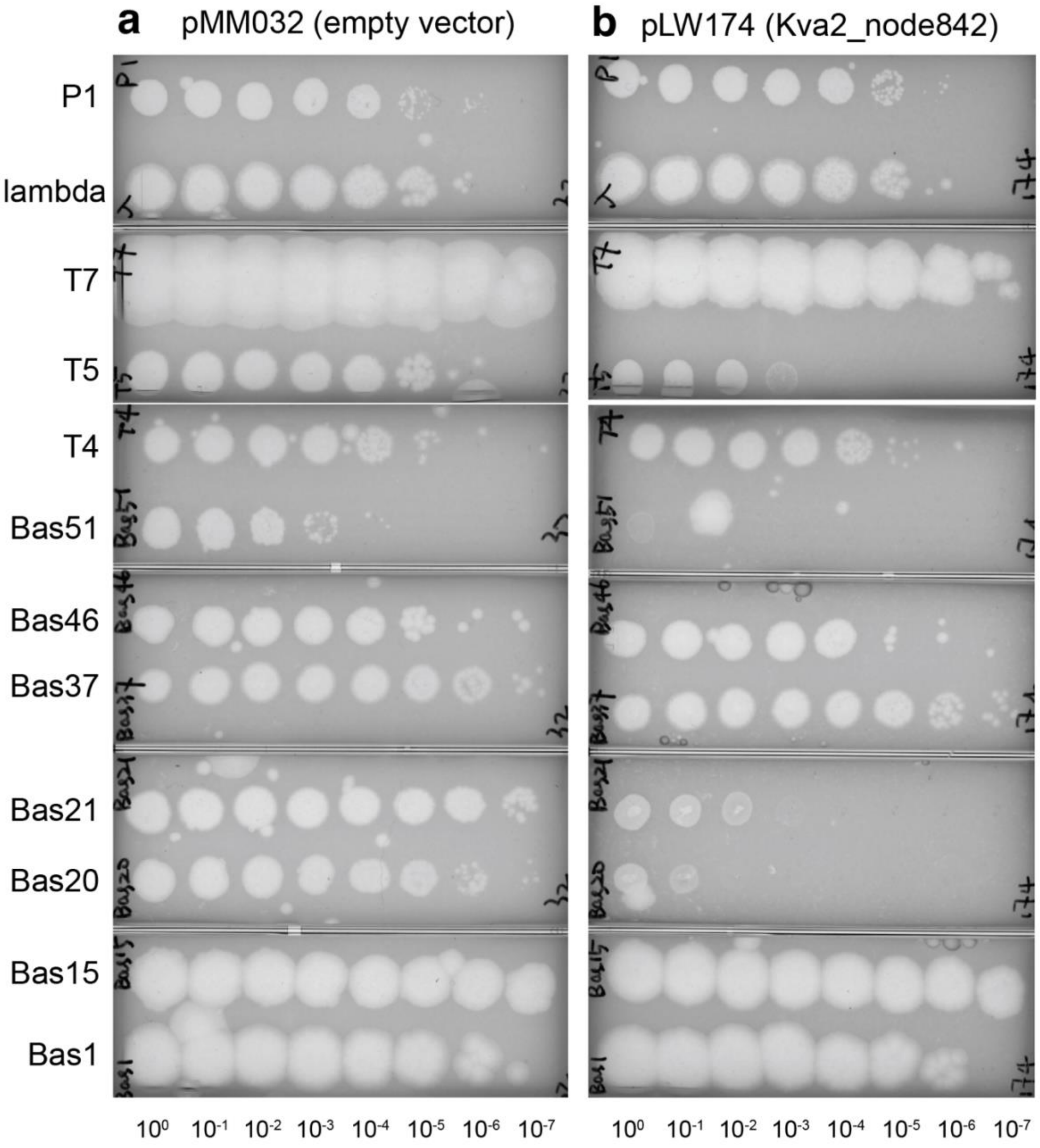


**Extended Data Fig. 1: Expanded phage panel testing retron-Kva2 defense. (a)** Spot dilution plaque assays of E. coli expressing an empty vector (pMM032) challenged with a diverse panel of bacteriophages: P1, λ, T7, T5, T4, Bas51, Bas46, Bas37, Bas21, Bas20, Bas15, and Bas1. **(b)** Plaque assays of *E. coli* expressing wild-type retron-Kva2 (pLW174) challenged with the identical phage panel. Ten-fold serial dilutions are spotted from $10^0$ to $10^{-7}$. Retron-Kva2 selectively restricts T5, Bas21, and Bas20.

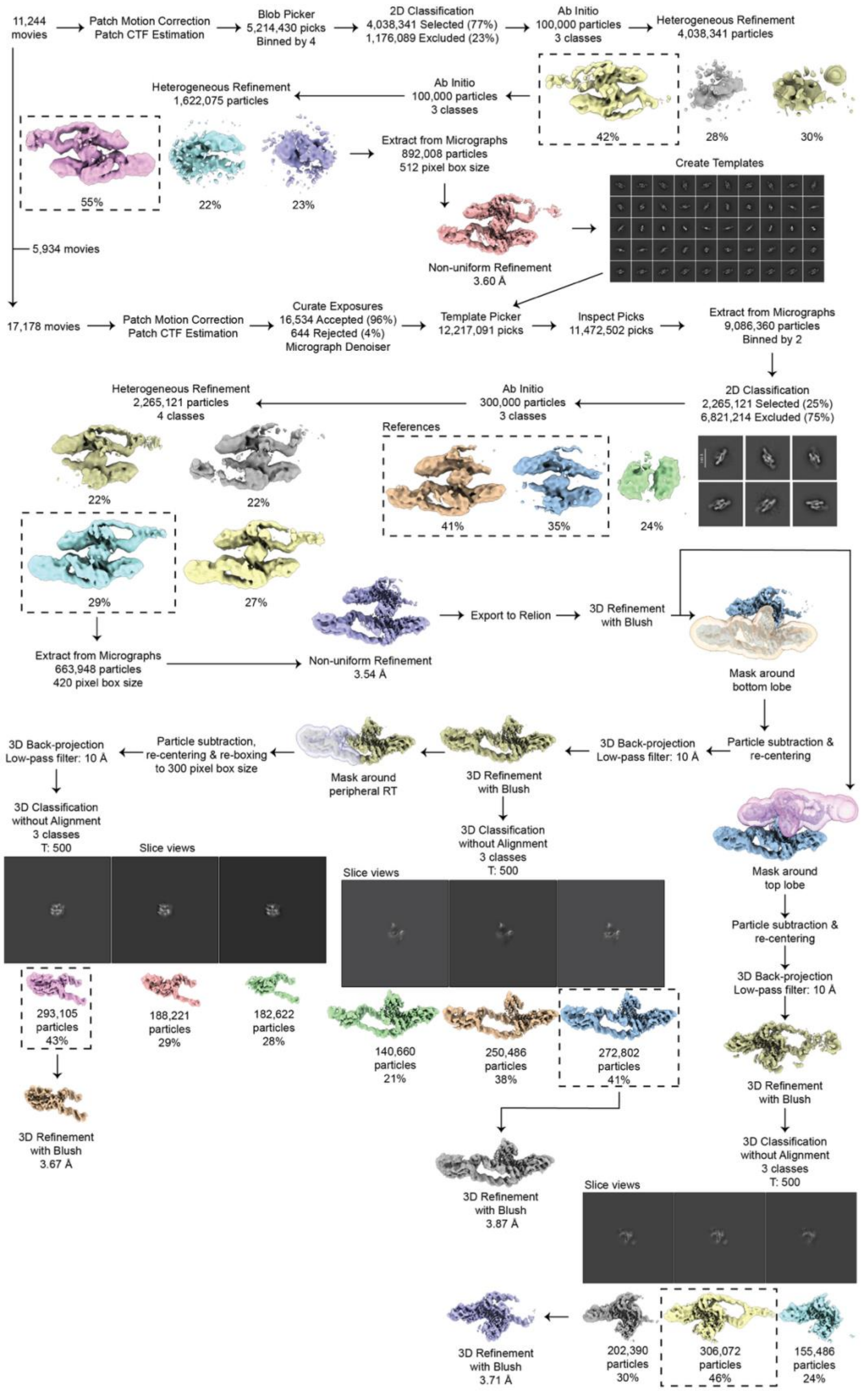
11,244 movies
Patch Motion Correction
Patch CTF Estimation
Blob Picker
5,214,430 picks
Binned by 4
2D Classification
4,038,341 Selected (77%)
1,176,089 Excluded (23%)
Ab Initio
100,000 particles
3 classes
Heterogeneous Refinement
4,038,341 particles
42%
28%
30%
Ab Initio
100,000 particles
3 classes
Heterogeneous Refinement
1,622,075 particles
55%
22%
23%
Extract from Micrographs
892,008 particles
512 pixel box size
Create Templates
Non-uniform Refinement
3.60 Å
5,934 movies
17,178 movies
Patch Motion Correction
Patch CTF Estimation
Curate Exposures
16,534 Accepted (96%)
644 Rejected (4%)
Micrograph Denoiser
Template Picker
12,217,091 picks
Inspect Picks
11,472,502 picks
Extract from Micrographs
9,086,360 particles
Binned by 2
2D Classification
2,265,121 Selected (25%)
6,821,214 Excluded (75%)
Ab Initio
300,000 particles
3 classes
References
41%
35%
24%
Heterogeneous Refinement
2,265,121 particles
4 classes
22%
22%
29%
27%
Extract from Micrographs
663,948 particles
420 pixel box size
Non-uniform Refinement
3.54 Å
Export to Relion
3D Refinement
with Blush
Mask around
bottom lobe
Particle subtraction &
re-centering
3D Back-projection
Low-pass filter: 10 Å
3D Refinement
with Blush
Mask around
peripheral RT
Particle subtraction,
re-centering & re-boxing
to 300 pixel box size
3D Back-projection
Low-pass filter: 10 Å
3D Classification
without Alignment
3 classes
T: 500
Slice views
293,105
particles
43%
188,221
particles
29%
182,622
particles
28%
3D Refinement
with Blush
3.67 Å
3D Classification
without Alignment
3 classes
T: 500
Slice views
140,660
particles
21%
250,486
particles
38%
272,802
particles
41%
3D Refinement
with Blush
3.87 Å
Mask around
top lobe
Particle subtraction &
re-centering
3D Back-projection
Low-pass filter: 10 Å
3D Refinement
with Blush
3D Classification
without Alignment
3 classes
T: 500
Slice views
202,390
particles
30%
306,072
particles
46%
155,486
particles
24%
3D Refinement
with Blush
3.71 Å

**Extended Data Fig. 2: Cryo-EM data processing workflow for the retron-Kva2 complex.** Schematic detailing the single-particle cryo-EM data processing pipeline. Initial motion correction and CTF estimation were performed in cryoSPARC Live[49]. Following template generation via blob picking, automated template-based picking in cryoSPARC (v5.0)[49] yielded an initial particle stack. After selective 2D classification to remove artifacts, particles were subjected to *ab initio* reconstruction and heterogeneous refinement. The best class was refined using non-uniform refinement and exported to RELION (v5.0) for 3D refinement with Blush[50]. To resolve distinct structural regions, masks were generated for the top and bottom lobes, which underwent independent particle subtraction, re-centering, 3D back-projection (10 Å low-pass filter), and 3D refinement with Blush. The peripheral RT was isolated from the bottom lobe, subtracted, re-centered, re-boxed (300 pixels), and low-pass filtered to 10 Å. Each localized component (top lobe, bottom lobe, and peripheral RT) was subjected to focused 3D classification without alignment (T=500). The resulting major classes were individually refined using 3D refinement with Blush.

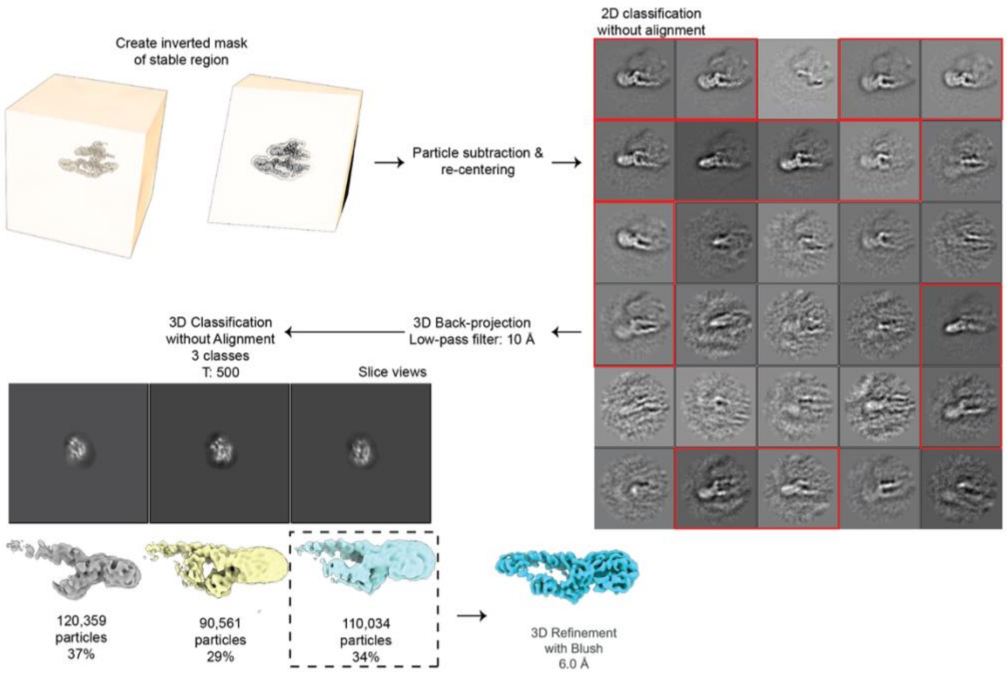


**Extended Data Fig. 3: Focused classification and local refinement of the retron-Kva2 flexible fourth RT.** An inverted mask of the stable structural core was generated to facilitate particle subtraction and re-centering. Subsequent 2D classification without alignment (red boxes indicate selected classes) and 3D classification without alignment isolated a distinct subpopulation of 110,034 particles displaying the flexible RT-msRNA-msdDNA. Final 3D refinement with Blush regularization yielded a 6.0 Å map of this dynamic subunit.

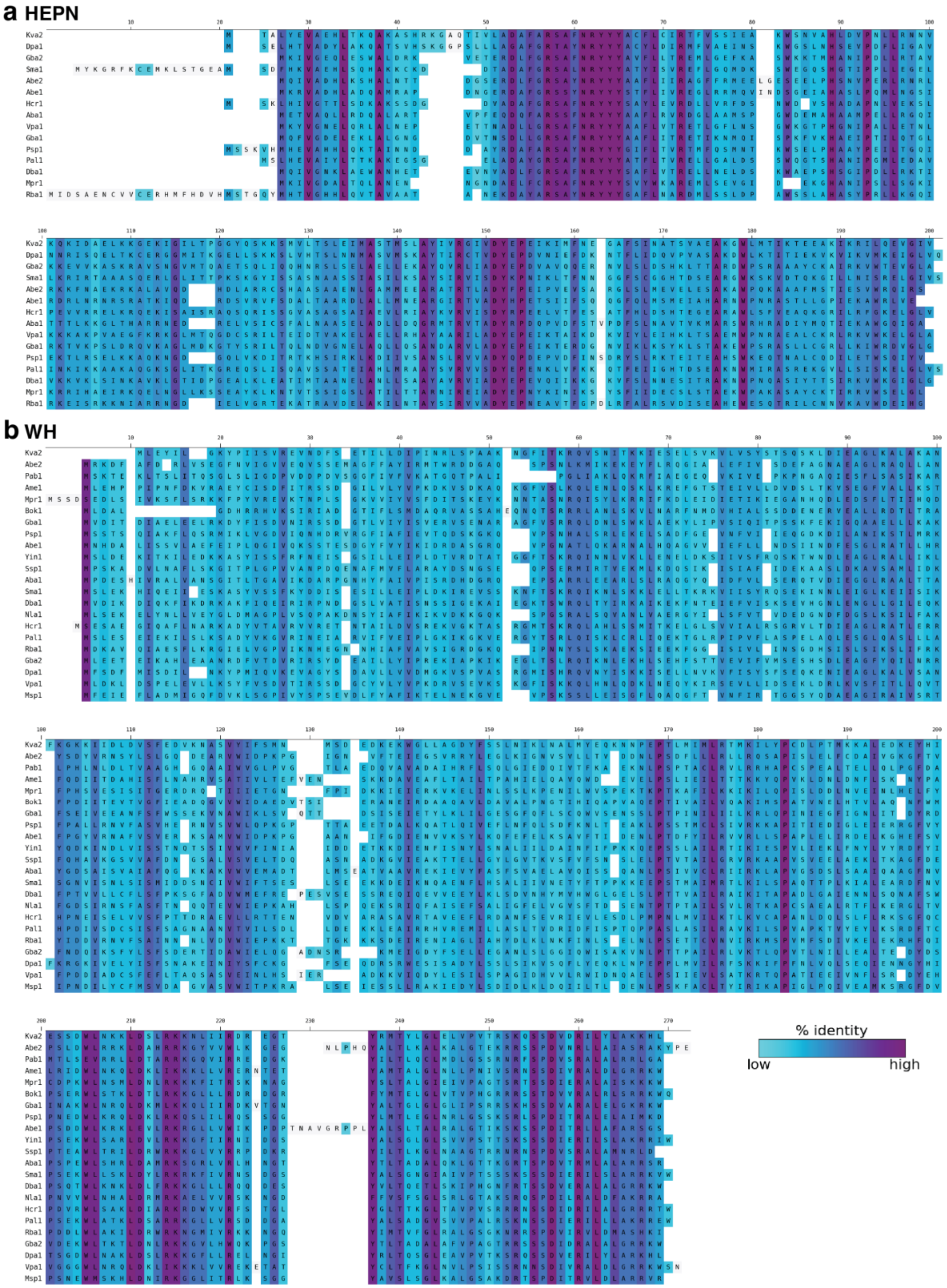
a HEPN
b WH
% identity
low
high

**c RT**

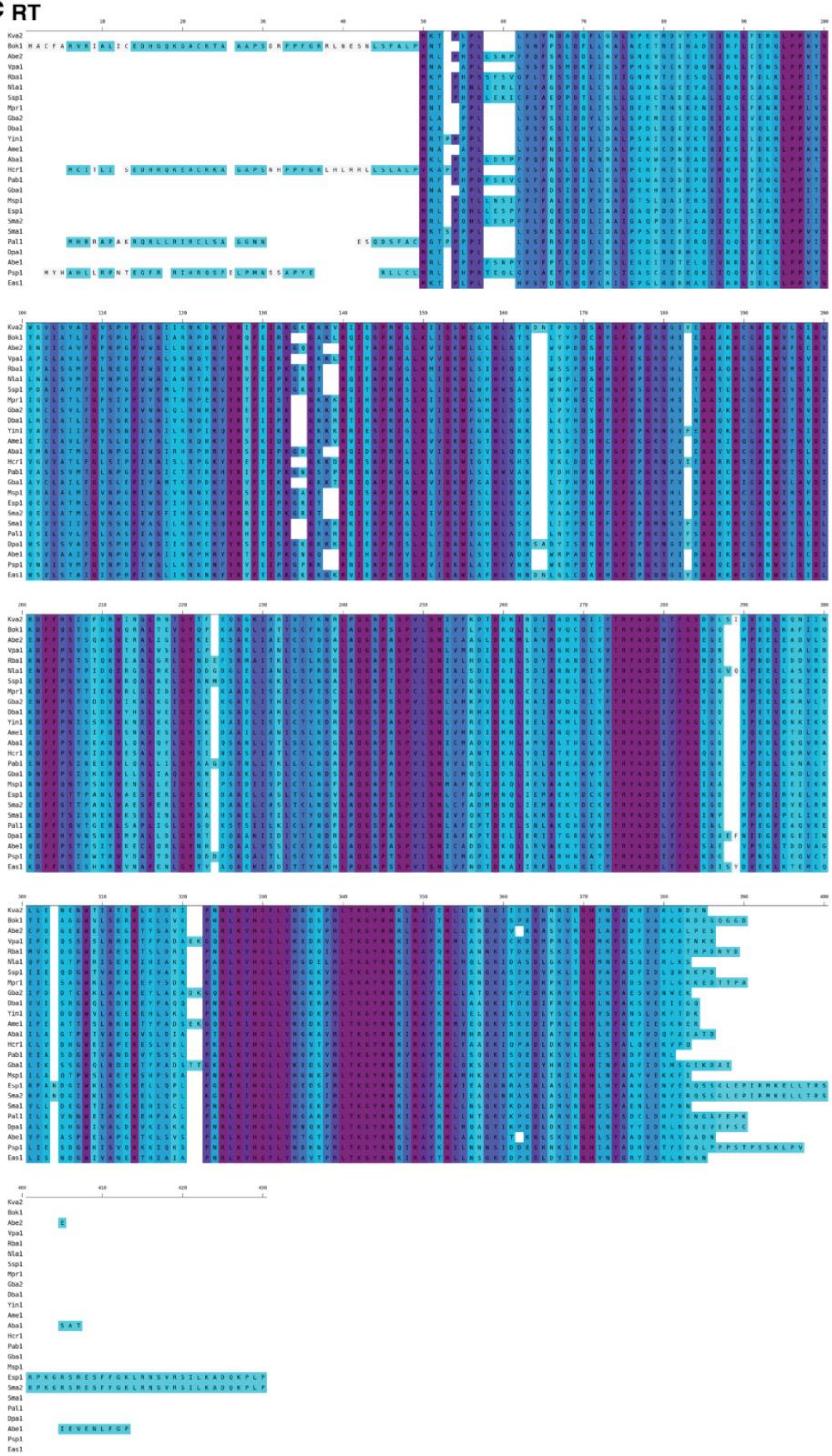

**d** ncRNA

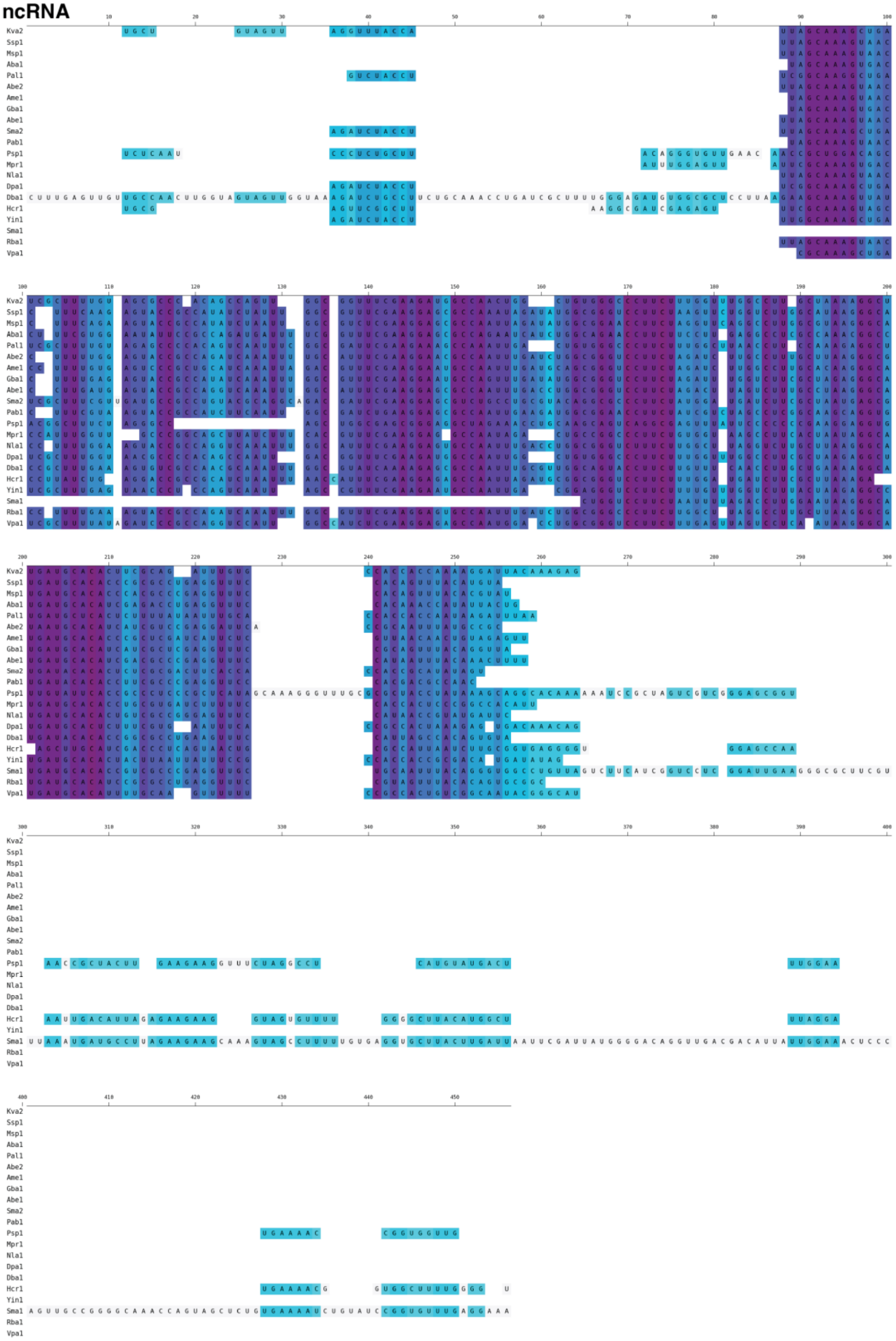

Kva2
Ssp1
Msp1
Aba1
Pal1
Abe2
Ame1
Gba1
Abe1
Sma2
Pab1
Psp1
Mpr1
Nla1
Dpa1
Dba1
Hcr1
Yin1
Sma1
Rba1
Vpa1

**Extended Data Fig. 4: Sequence conservation across type IX retron components.** Multiple sequence alignments of the retron-Kva2 operon components against type IX retron defense systems predicted by Mestre et al.[12]. **(a)** HEPN effector protein. **(b)** WH effector protein. **(c)** RT protein **(d)** ncRNA sequences. Residues and nucleotides are colored by percent identity (dark purple indicates high conservation; light blue indicates low conservation).

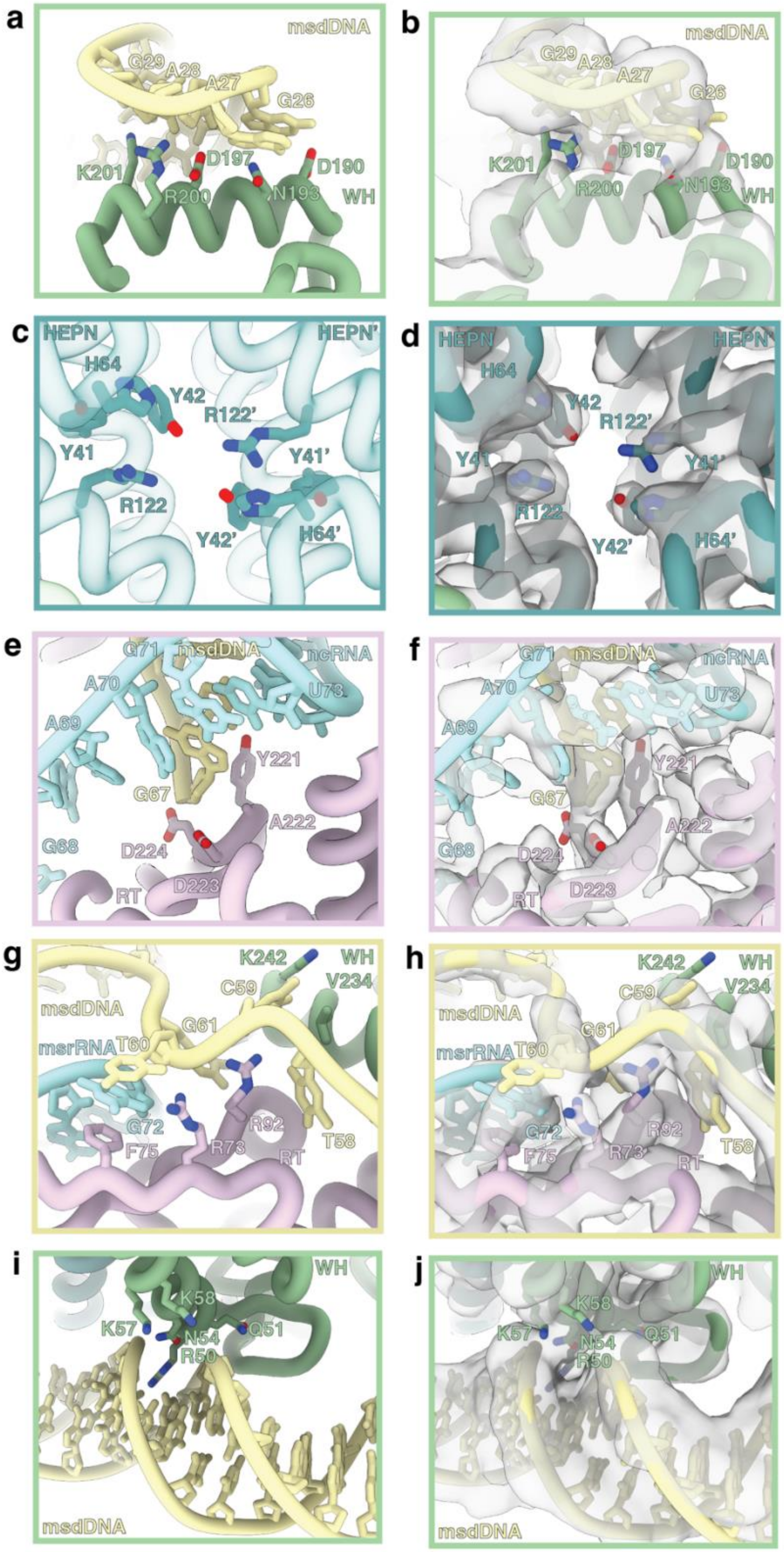

a
msdDNA
G29
A28
A27
G26
K201
D197
D190
R200
N193
WH
b
msdDNA
G29
A28
A27
G26
K201
D197
D190
R200
N193
WH
c
HEPN
HEPN'
H64
Y42
R122'
Y41
Y41'
R122
Y42'
H64'
d
HEPN
HEPN'
H64
Y42
R122'
Y41
Y41'
R122
Y42'
H64'
e
G71
msdDNA
ncRNA
A70
U73
A69
Y221
G67
A222
G68
D224
D223
RT
f
G71
msdDNA
ncRNA
A70
U73
A69
Y221
G67
A222
G68
D224
D223
RT
g
K242
WH
C59
V234
msdDNA
G61
msrRNA
T60
G72
R92
F75
R73
RT
T58
h
K242
WH
C59
V234
msdDNA
G61
msrRNA
T60
G72
R92
F75
R73
RT
T58
i
WH
K58
K57
Q51
N54
R50
msdDNA
j
WH
K58
K57
Q51
N54
R50
msdDNA

**Extended Data Fig. 5: Models and cryo-EM density maps of key functional interfaces within the retron-Kva2 complex.** Atomic models (left panels) and corresponding cryo-EM density maps overlaid on the models (right panels) highlighting critical structural regions. **(a, b)** The WH-msdDNA interface, depicting interactions between WH residues D190, N193, D197, R200, and K201 with msdDNA bases. **(c, d)** The symmetric HEPN dimer active site, showing the catalytic residues H64 and R122 alongside substrate-positioning residues Y41 and Y42 from each monomer. **(e, f)** The RT active site cradling the processed msdDNA, featuring the conserved YADD motif (Y221, A222, D223, D224). **(g, h)** Proximal WH-msdDNA and msrRNA contacts near the RT domain. **(i, j)** Distal WH minor groove interactions along the msdDNA stem loop.

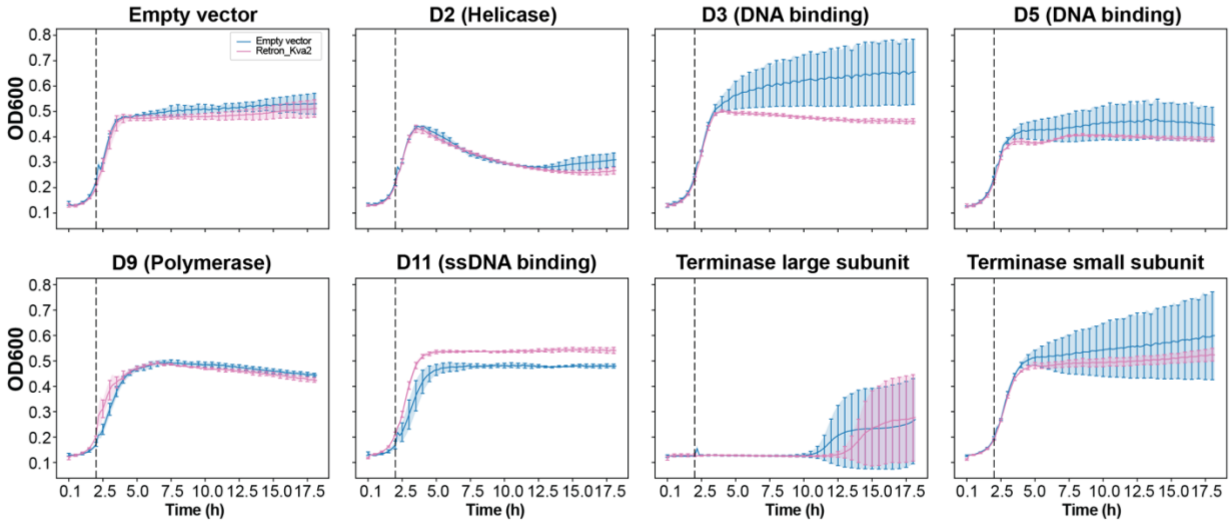


**Extended Data Fig. 6: Co-expression growth curves of retron-Kva2 and candidate phage triggers.** T5 genes discovered from sequencing of the T5 phage escapees were cloned into individual plasmids under arabinose-inducible promoters. The candidate phage triggers were co-expressed either with retron-Kva2 or empty vector. Their growth was monitored for 18 hours.

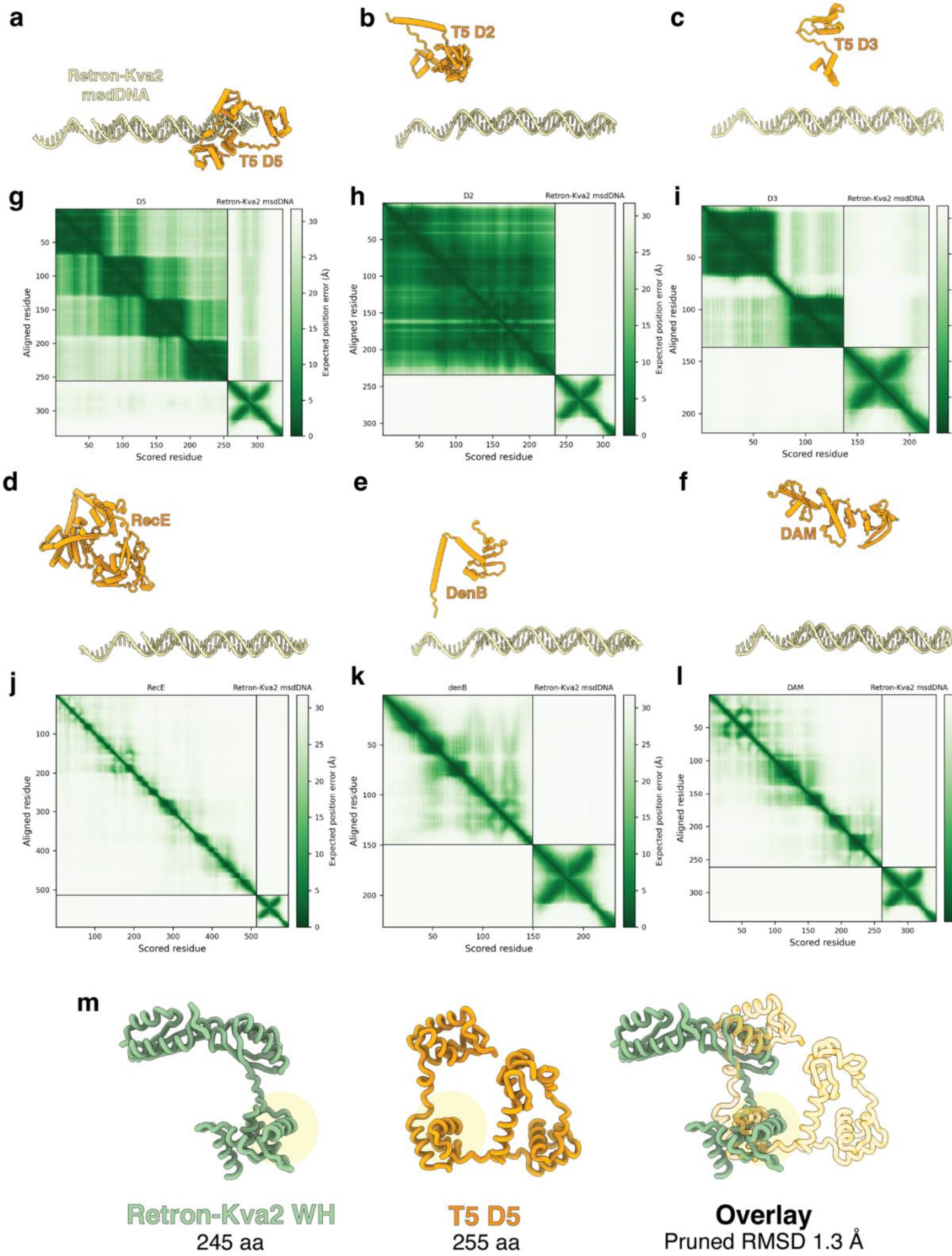
a
Retron-Kva2
msdDNA
T5 D5
b
T5 D2
c
T5 D3
g
D5
Retron-Kva2 msdDNA
Aligned residue
Scored residue
Expected position error (Å)
h
D2
Retron-Kva2 msdDNA
Aligned residue
Scored residue
Expected position error (Å)
i
D3
Retron-Kva2 msdDNA
Aligned residue
Scored residue
Expected position error (Å)
d
RecE
e
DenB
f
DAM
j
RecE
Retron-Kva2 msdDNA
Aligned residue
Scored residue
Expected position error (Å)
k
denB
Retron-Kva2 msdDNA
Aligned residue
Scored residue
Expected position error (Å)
l
DAM
Retron-Kva2 msdDNA
Aligned residue
Scored residue
Expected position error (Å)
m
Retron-Kva2 WH
245 aa
T5 D5
255 aa
Overlay
Pruned RMSD 1.3 Å

**Extended Data Fig. 7: AlphaFold3 structural predictions and PAE plots of candidate phage triggers.** (a-f) AlphaFold3[26] structure predictions of candidate phage proteins docked against the retron-Kva2 msdDNA stem loop, testing T5 proteins D5 **(a)**, D2 **(b)**, and D3 **(c)**, alongside proteins RecE **(d)**, DenB **(e)**, and DAM **(f)**. **(g-l)** Corresponding Predicted Aligned Error (PAE) plots for each predicted complex. A strong interaction signal (low off-diagonal expected position error) is observed for the D5-msdDNA interaction **(g)**. **(m)** Structural overlay of the experimentally determined retron-Kva2 WH domain (green) and the predicted T5 D5 structure (orange), demonstrating topological mimicry of the HTH fold with a pruned RMSD of 1.3 Å.

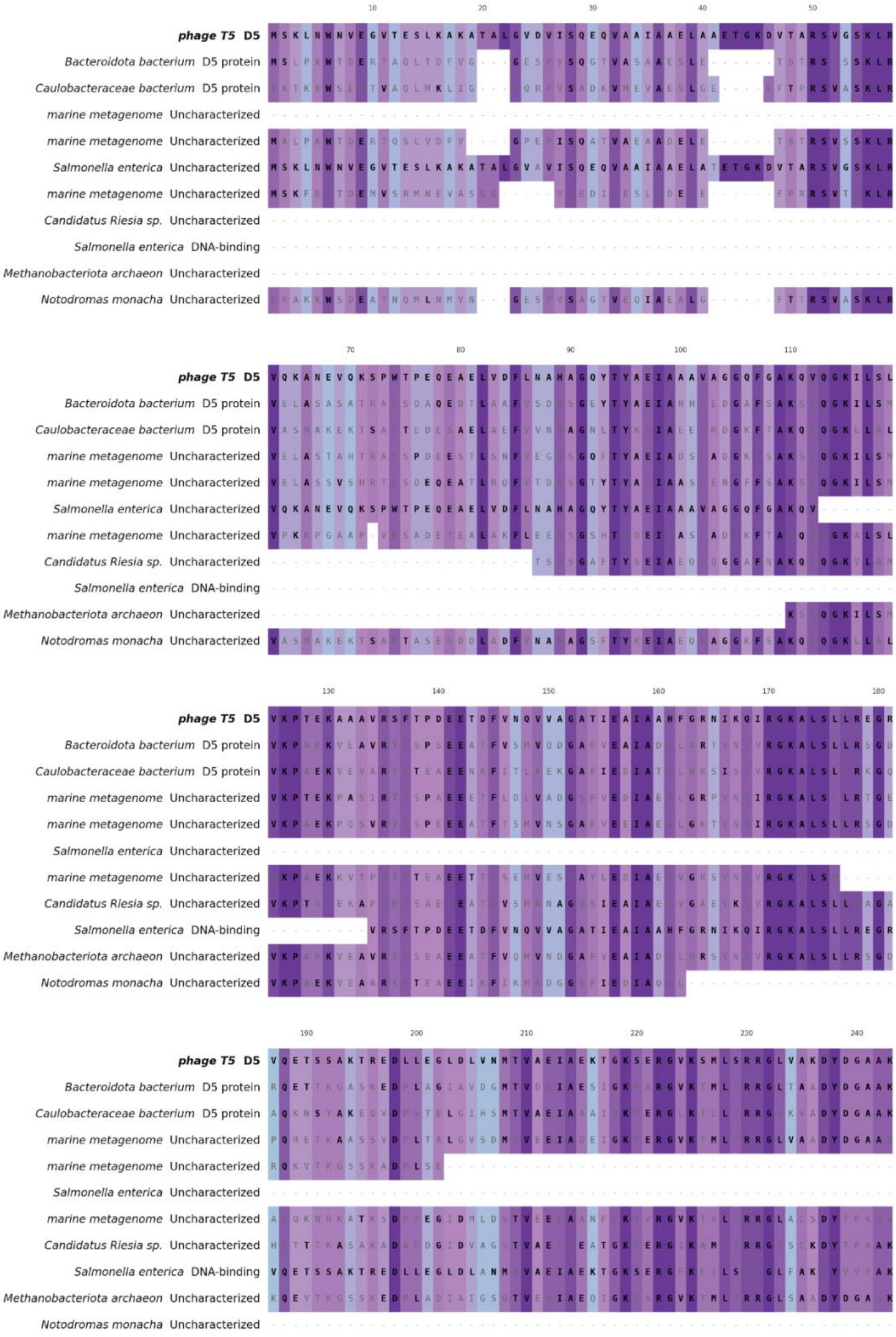
phage T5 D5
Bacteroidota bacterium D5 protein
Caulobacteraceae bacterium D5 protein
marine metagenome Uncharacterized
marine metagenome Uncharacterized
Salmonella enterica Uncharacterized
marine metagenome Uncharacterized
Candidatus Riesia sp. Uncharacterized
Salmonella enterica DNA-binding
Methanobacteriota archaeon Uncharacterized
Notodromas monacha Uncharacterized

**Extended Data Fig. 8: Multiple sequence alignment of phage T5 D5 and diverse structural homologs.** Sequence alignment of the T5 D5 trigger protein against structural homologs identified from FoldSeek[28]. Residues are colored according to percent identity.

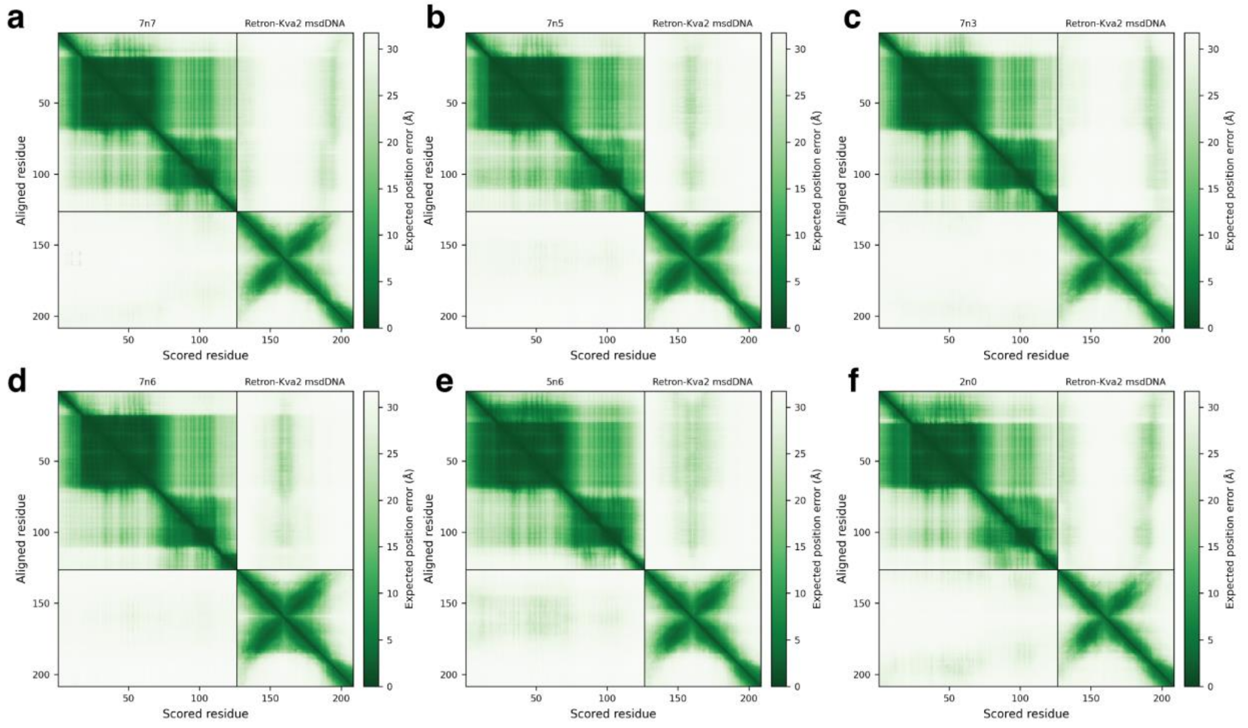


**Extended Data Fig. 9: Confidence metrics for *de novo* designed synthetic triggers.** PAE plots generated by AlphaFold3[26] for the six computationally designed synthetic triggers with retron-Kva2 msdDNA: **(a)** 7n7, **(b)** 7n5, **(c)** 7n3, **(d)** 7n6, **(e)** 5n6, and **(f)** 2n0.

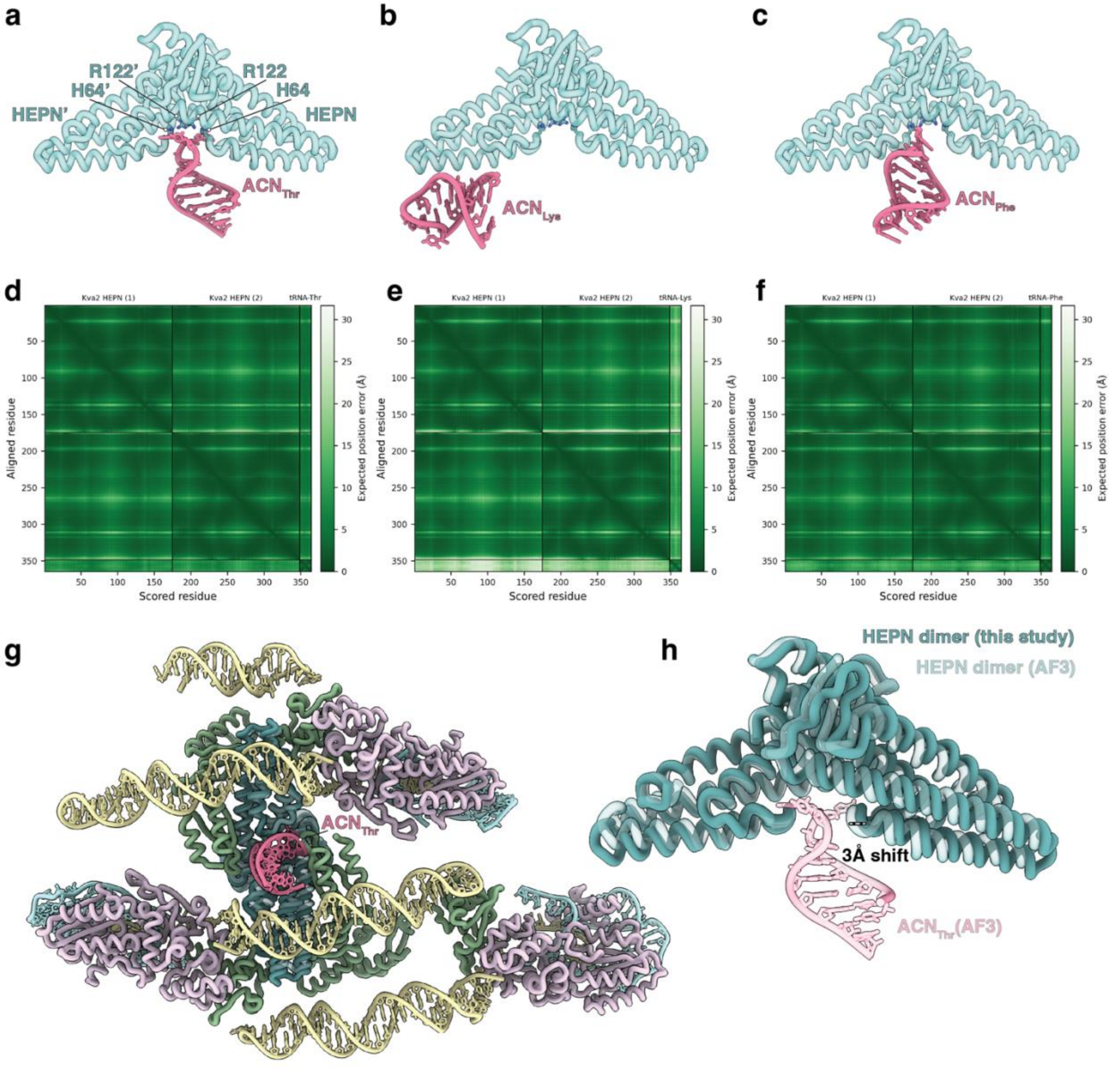


**Extended Data Fig. 10: HEPN–tRNA engagement and the structural basis of retron-Kva2 activation. (a–c)** AlphaFold3[26] structural predictions of the retron-Kva2 HEPN homodimer (cyan) evaluated against candidate tRNA anticodon stem loops (pink): **(a)** threonine ($ACN_{Thr}$), **(b)** lysine ($ACN_{Lys}$), and **(c)** phenylalanine ($ACN_{Phe}$). The $ACN_{Thr}$ stem loop is uniquely predicted to dock in a catalytically competent orientation within the bipartite active site cleft formed by conserved residues H64, R122, H64′, and R122′. **(d–f)** Corresponding PAE plots for the structural models in **a–c**, displaying the expected position error (Å) between the two HEPN monomers and their respective tRNA ACN targets. (g) The predicted HEPN–$ACN_{Thr}$ complex overlaid with the experimentally determined retron-Kva2 higher-order architecture, demonstrating the location of tRNA substrate entry. **(h)** Structural superposition of the autoinhibited HEPN dimer determined by cryo-EM in this study (opaque) and the AlphaFold3-predicted active HEPN dimer bound to $ACN_{Thr}$ (transparent). Substrate engagement correlates with a predicted 3 Å conformational shift at the active site, revealing the structural rearrangement necessary to unleash collateral tRNase activity.

**Extended Data Table 1: Cryo-EM data collection, refinement and validation statistics**

| | Retron-Kva2 (PDB 36HT, EMDB-77585) |
|---|---|
| **Data collection and processing** | |
| Magnification | 36,000 |
| Voltage (kV) | 200 |
| Electron exposure (e–/Å²) | 50 |
| Defocus range (μm) | 1-3 |
| Pixel size (Å) | 1.14 |
| Symmetry imposed | C1 |
| Initial particle images (no.) | 9,086,360 |
| Final particle images (no.) | 663,948 |
| Map resolution (Å) | 3.7 |
| *FSC threshold* | 0.143 |
| Map resolution range (Å) | 2.5-7 Å |
| **Refinement** | |
| Initial model used (PDB code) | |
| Model resolution (Å) | 2.5 |
| *FSC threshold* | 0.143 |
| Model resolution range (Å) | 1.7-4.0 |
| Map sharpening B factor (Å²) | |
| Model composition | |
| *Non-hydrogen atoms* | 22,778 |
| *Protein residues* | 1,955 |
| *Nucleotide residues* | 339 |
| *Ligands* | 0 |
| B factors (Å²) | |
| *Protein* | 107.53 |
| *Nucleotide* | 156.96 |
| *Ligand* | |
| R.m.s. deviations | |
| *Bond lengths (Å)* | 0.006 |
| *Bond angles (°)* | 1.168 |
| Validation | |
| *MolProbity score* | 1.35 |
| *Clashscore* | 4.56 |
| *Poor rotamers (%)* | 0.68 |
| Ramachandran plot | |
| *Favored (%)* | 97.36 |
| *Allowed (%)* | 2.64 |
| *Disallowed (%)* | 0.00 |

**Extended Data Table 2: Mutations found in T5 phage escapees**

| Phage escapees | Mutated gene product | Mutation location in T5 genome | Mutation | Mutation on protein |
|---|---|---|---|---|
| T5.1 | D2, helicase | 63347 | G→C | E76Q |
| T5.2 | D3, DNA binding protein | 64261 | G→C | E30Q |
| T5.2 | D3, DNA binding protein | 64427 | G→A | G85D |
| T5.1 | D5, DNA binding protein | 67467 | G→A | G53D |
| T5.1 | D5, DNA binding protein | 67530-67531 | GG→AA | W74* |
| T5.1 | D5, DNA binding protein | 67754 | G→A | V149I |
| T5.1 | D5, DNA binding protein | 68004 | G→A | G232D |
| T5.1 | D9, polymerase | 71785 | A→C | Y230S |
| T5.1 | D9, polymerase | 71790 | T→G | H231Q |
| T5.1 | D9, polymerase | 71789 | C→A | L232M |
| T5.1 | D9, polymerase | 71794 | G→T | G233V |
| T5.2 | D11, ssDNA binding protein | 76157 | G→C | E53Q |
| T5.2 | D11, ssDNA binding protein | 76193 | G→A | V65I |
| T5.2 | D11, ssDNA binding protein | 76200 | G→A | G67D |
| T5.2 | D11, ssDNA binding protein | 76208 | G→A | D70N |
| T5.2 | D11, ssDNA binding protein | 76227 | G→A | G76D |
| T5.2 | D11, ssDNA binding protein | 76265 | G→A | V89I |
| T5.1 | Terminase large subunit | 107247 | G→A | R230C |
| T5.1 | Terminase large subunit | 107249 | G→A | A229V |
| T5.1 | Terminase large subunit | 107594 | G→A | T114I |
| T5.1 | Terminase large subunit | 107772 | G→A | R55C |
| T5.1 | Terminase large subunit | 107775 | G→A | H54Y |
| T5.2 | Terminase small subunit | 107984 | G→A | Q145* |
| T5.2 | Terminase small subunit | 108022 | G→A | T132I |
